# The influence of N-methylation on the ansamers of an amatoxin: Gly5Sar-amanullin

**DOI:** 10.1101/2022.12.21.521444

**Authors:** Marius T. Wenz, Simone Kosol, Guiyang Yao, Roderich D. Süssmuth, Bettina G. Keller

**Affiliations:** Freie Universität Berlin, Department of Biology, Chemistry and Pharmacy, Arnimallee 22, 14195 Berlin, Germany; Technische Universität Berlin, Institute for Chemistry, Strasse des 17. Juni 124, 10623 Berlin, Germany; Fudan University, Center for Innovative Drug Discovery, Greater Bay Area Institute of Precision Medicine (Guangzhou), School of Life Sciences, PR China

## Abstract

Amatoxins are strong inhibitors of RNA polymerase II, and cause cell death. Because of their cytotoxicity they are candidates for anti-cancer drugs, and understanding their structure-activity relationship is crucial. Amatoxins have a rigid bicyclic scaffold which consists of a cyclic octapeptide bridged by cysteine and tryptophan side chain forming a tryptathionine bridge. Here we show the influence of the N-methylation on the amatoxin scaffold by studying Gly5Sar-amanullin with MD simulations and NMR experiments. Since we have shown recently that the amatoxin scaffold allows for two isomeric forms (ansamers), we studied both isomers of Gly5Sar-amanullin. We found that both isomers of Gly5Sar-amanullin form two long-living conformations which is unusual for amatoxins, and that they are differently affected by the N-methylation. The natural Gly5Sar-amanullin forfeits the hydrogen bonds to Gly5 due to the N-methylation, which is expected from existing crystal structures for alpha-amanitin. Our results however indicate that this does not cause more flexibility due to a shift in the hydrogen bond pattern. In the unnatural isomer, we observe an interesting cis-trans-isomerisation of the backbone angles in Trp4 and Gly7, which is enabled by the N-methylation. We expect that our perspective on the effect of N-methylation in amatoxins could be a starting point for further SAR-studies which are urgently needed for the design of better anti-cancer agents.

## Introduction

Amatoxins are a family of “utterly powerful poisons” ^1^ that occur in several poisonous mushrooms, most-notably the death cap mushroom (*Amanita phalloides*).^2–6^ They consist of eight amino acids with the general sequence Asn-Pro-Ile-Trp-Gly-Ile-Gly-Cys which are joined in a cyclic peptide chain (macrolactam). Additionally, amatoxins have a covalent bridge between cysteine and tryptophan (tryptathionine bridge), such that their overall structure is a bicyclic octapeptide. The various members for the amatoxin family differ in the hydroxylation and amination of residues Asp1, Pro2, Ile3, and Trp4.^2^ The most prominent member of this family is *α*-amanitin.^7^

Amatoxins are stable at temperatures beyond 100 °C and thus cannot be destroyed by cooking the poisonous mushrooms. They are water-soluble which facilitates their oral uptake. Finally, their exceptional structure makes them resistant against stomach acid or digestion by proteases.^2,3,8^ They act by inhibiting RNA polymerase II, thereby shutting down protein synthesis and cell metabolism.^9–12^ By interacting with cell membranes, amatoxins can also be responsible for cell lysis and misplaced cell organelles.^2^ Amatoxins can affect all organs, but their effect on liver and heart is most destructive, and a fatally poisoned animal or human usually dies of liver or heart failure.^13–15^ No definitive antidote is available, and, in case of liver failure, a liver transplant is the only viable treatment.^5,15^

Because amatoxins are so extraordinarily cytotoxic, they are ideal candidates for targeted anti-cancer therapy.^16,17^ In this approach, one develops an antibody that binds to an antigen on the surface of the cancer cell and couples it to a potent cytoxin. After uptake by the cancer cell, the cytoxin is cleaved from the antibody, and the released cytotoxin kills the cancer cell.

Even though amatoxins are very promising payloads for antibody-drug conjugates, ^16,18^ they could not be used in cancer research until recently, because their synthesis is extremely challenging. In mushrooms, amatoxins are ribosomally synthesised,^4,19^ but obtaining amatoxins in sufficient quantity and quality from mushroom farms proved to be impossible.^16,20,21^ Until today, there have only been four reports on the successful synthesis of *α*-amanitin, all of which were published in the last five years.^20,22–24^ We recently proposed a very efficient synthesis for amatoxin,^24^ which allowed us to build an amatoxin library to systematically study their structure-activity relationship.^25^

Each amatoxin can in principle occur as two different isomers, which differ in the position of the tryptathionine bridge relative to the macrolactam ring. Here, the two sides of the macrolactam ring are defined by the direction of peptide chain: NH → C_*α*_ → CO. When looking down on the macrolactam ring and the peptide chain proceeds clock-wise, one looks at the “upper side” of the ring. So far, only the isomer, in which the bridge is located above the macrolactam ring has been isolated from mushrooms.

The existence of the second isomer, in which the tryptathionine bridge is below the macrolactam ring, was highly debated in literature. ^26–28^ However, recently we were able to synthesise, isolate and characterise both isomers of the amatoxin amanullin.^29^ We also proposed the term ‘ansamers’ for this particular isomerism, where *P*_ansa_ is the natural isomer with the bridge above the macrolactam ring, and *M*_ansa_ is the newly synthesized non-natural isomer with the bridge below the macrolactam ring. Formally, the two ansamers can be interconverted by rotating the tryptathionine bridge around the (imaginary) axis between the two bridgeheads, but this rotation is hindered by the macrolactam ring. Consequently, the ansamers are in fact two separate substances, which differ in their structure and physicochemical properties. For example, the *M* -ansamer of amanullin has a different intramolecular hydrogen-bond pattern which makes it more compact and more hydrophobic than the *P* - ansamer.

Monocyclic peptides often exhibit a variety of conformations which differ in their physicochemical properties and interconvert on relatively slow timescales, often microseconds or slower^30–33^ However, the interconversion is usually not slow enough to separate the conformations by chromatography or similar techniques. Yet, the conformational dynamics are crucial for understanding and tuning the structure-activity relationship of monocyclic peptides.^28,30,31,34,35^ A classic example is cyclosporine which can interconvert between a hydrobphilic conformation which facilitates oral uptake, and a hydrophobic conformation which facilitates membrane permeation.^36–39^ With this in mind, two questions arise: (1) Can amatoxins exhibit several conformations, or does the tryptathionine bridge enforce one dominant conformation? (2) How do the conformations differ in the *P* -ansamer and in the *M* -ansamer? In our previous study, we studied amanullin which differs from *α*-amanitin in the hydroxylation of the side chains in position 4 and 6. At position 4, it contains a normal tryptophan instead of the hydroxylated 6-OH-tryptophan (OHTrp), and at position 6, it contains a normal isoleucine instead of the twice hydroxylated (3*R*,4*R*)-4,5-dihydroxyisoleucine (DHIle). Spectroscopic data indicated that amanullin only forms a single dominant backbone conformation in each of the ansamers of amanullin (with two conformations of the Ile3-ethyl group in the crystal structure of the *M* -ansamer).^29^ The conformation of the *P* -ansamer is stabilized by a strong hydrogen bond from the amide of Gly5 to the carbonyl oxygen of Asn1. This hydrogen bond is known to contribute to the overall rigidity of the amatoxin scaffold.^29,40,41^

Here, we investigate the amatoxin Gly5Sar-amanullin, in which the amide of Gly5 has been methylated (sarcosine), thereby prohibiting the stabilizing hydrogen bond. N-methylation is a well-established strategy to modify the structure and dynamics of monocyclic peptides.^42,43^ We combine classical molecular dynamics (MD) simulations of the *P* -ansamer and *M* -ansamer of Gly5Sar-amanullin with NMR experiments of the *M* -ansamer to identify the stable conformations of Gly5Sar-amanullin. We show that the methylation has a very different affect on the *P* -ansamer than on the *M* -ansamer.

## Results and discussion

### Gly5Sar-amanullin

The conformation of *α*-amanitin and *P*_ansa_-amanullin is stabilized by a hydrogen bond between the amide hydrogen of Gly5 and the backbone carbonylgroup of Asn1 (Figure 1.c).^29^ This hydrogen bond is also found in the crystal structures of other amatoxins.^44^ To investigate the effects of the loss of this hydrogen bond on the structure and conformational stability, we synthesized a derivative with the N-methylated amino acid sarcosine (three-letter code: Sar) instead of glycine 5 (Figure 1.b).

**Figure 1:**
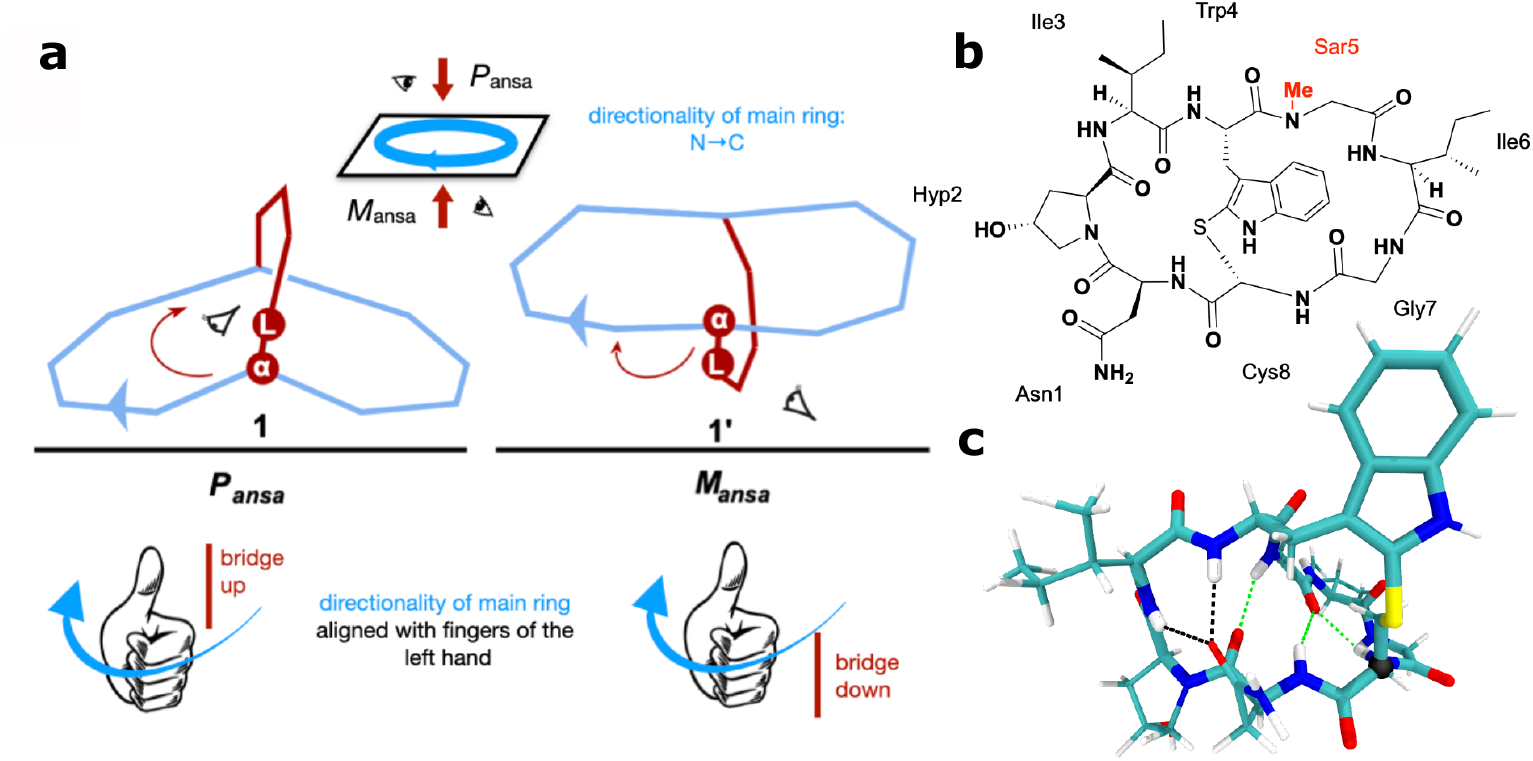
a) Definition of the ansamers *P*_ansa_ and *M*_ansa_ (Figure adapted from Ref. 29). b) Chemical structure of Gly5Sar-amanullin. c) Crystal structure of *P*_ansa_-amanullin (CCDC deposition numbers: 1128063^41^). The structure is coloured according to atom type: *cyan:* carbon; *white:* hydrogen; *red:* oxygen; *blue:* nitrogen; *yellow:* sulphur. The C_*α*_-atom of Cys8 is highlighted as black sphere. Hydrogen bonds are shown as dotted lines and coloured green if Gly5 is involved.

Ansamers arise in bicyclic structures in which at least one of the cycles has a clear directionality (main ring). For amatoxins, this ring is the peptide ring (macrolactam), where the direction is defined by the sequence N → C _*α*_ → carbonyl-C. Because of this directionality, the main ring has two distinct sides, and the position of the second cycle with respect to the main ring defines two distinct isomers. In our previous study,^29^ we suggested the term ansamers for this type of isomerism, with *P* -ansamer denoting one option and *M* -ansamer the other. To assign the stereochemical descriptor (*M* or *P* configuration), the CIP (CahnIngold-Prelog) rules can be followed.^29^ One proceeds as follows (Figure 1.a):

1. Identify the main ring and its directionality.
2. Identify the bridge-head atom *α* where the second cycle branches from the main ring.
3. Identify the leading atom *L* in the second cycle.
4. If looking along direction *L* → *α*,

- the main ring proceeds clockwise, the structure is a *P* -ansamer
- the main ring proceeds anti-clockwise, the structure is a *M* -ansamer.

Because, the second cycle always has two bridgeheads, one has two choices in step 2, but both choices lead to the same result. In amatoxins, the second cycle is the tryptathionine bridge, and the *P* -ansamer is often described as “bridge up”, whereas the *M* -ansamer is described as “bridge down”. This becomes understandable if one considers an equivalent procedure to identify the ansamers which uses a “left-hand rule” (Figure 1.a)

1. Identify the main ring and its directionality.
2. Curve the fingers of the left-hand in the directionality of the main ring. The direction of the thumb defined “up”.
3. If the second cycle and the thumb are located

- on the same side, the structure is a *P* -ansamer
- on different sides, the structure is a *M* -ansamer.

So far, only the *P* -ansamers of amatoxins have been isolated from mushrooms,^16^ but in the laboratory we could also synthesise *M* -ansamers.^24,29^

We synthesised Gly5Sar-amanullin (Figure 1.b) using our previously developed strategy:^24,29^ the iodine-mediated tryptathionine bridge formation and the bicycle ring-closure between Hyp2 and Ile3. We verified the product by HPLC-HRMS, ^1^H-NMR spectroscopy and UV-VIS spectroscopy. With our current protocol, we obtained exclusively one of the two possible ansamers, and the UV-VIS spectrum, CD spectrum and NMR NOE data suggest that it is the *M* -ansamer of Gly5Sar-amanullin (see Supplementary Information, sections 1 and 2).

### The *M* -ansamer has two populations in slow exchange

The ^1^H-NMR spectrum of the *M* -ansamer of Gly5Sar-amanullin showed two distinct peaks for the amine of the tryptophan side chain, suggesting two different conformers for the *M* -ansamer: 1_*M*_ and 2_*M*_ (Figure 2.a). From the integral under the two peaks in the 1D spectrum, we estimated the relative populations to be 37% for 1_*M*_ and 62% for 2_*M*_ at 298 K. The spectrum also shows a very small third peak corresponding to less than 1% of the population. The peaks in the ^1^H-NMR spectrum correspond to only a single chemical species, suggesting that 1_*M*_ and 2_*M*_ represent different conformers of the *M* -ansamer (Note that the subscript *M* in 1_*M*_ and 2_*M*_ stands for *M* -ansamer and not for magnetization.)

**Figure 2:**
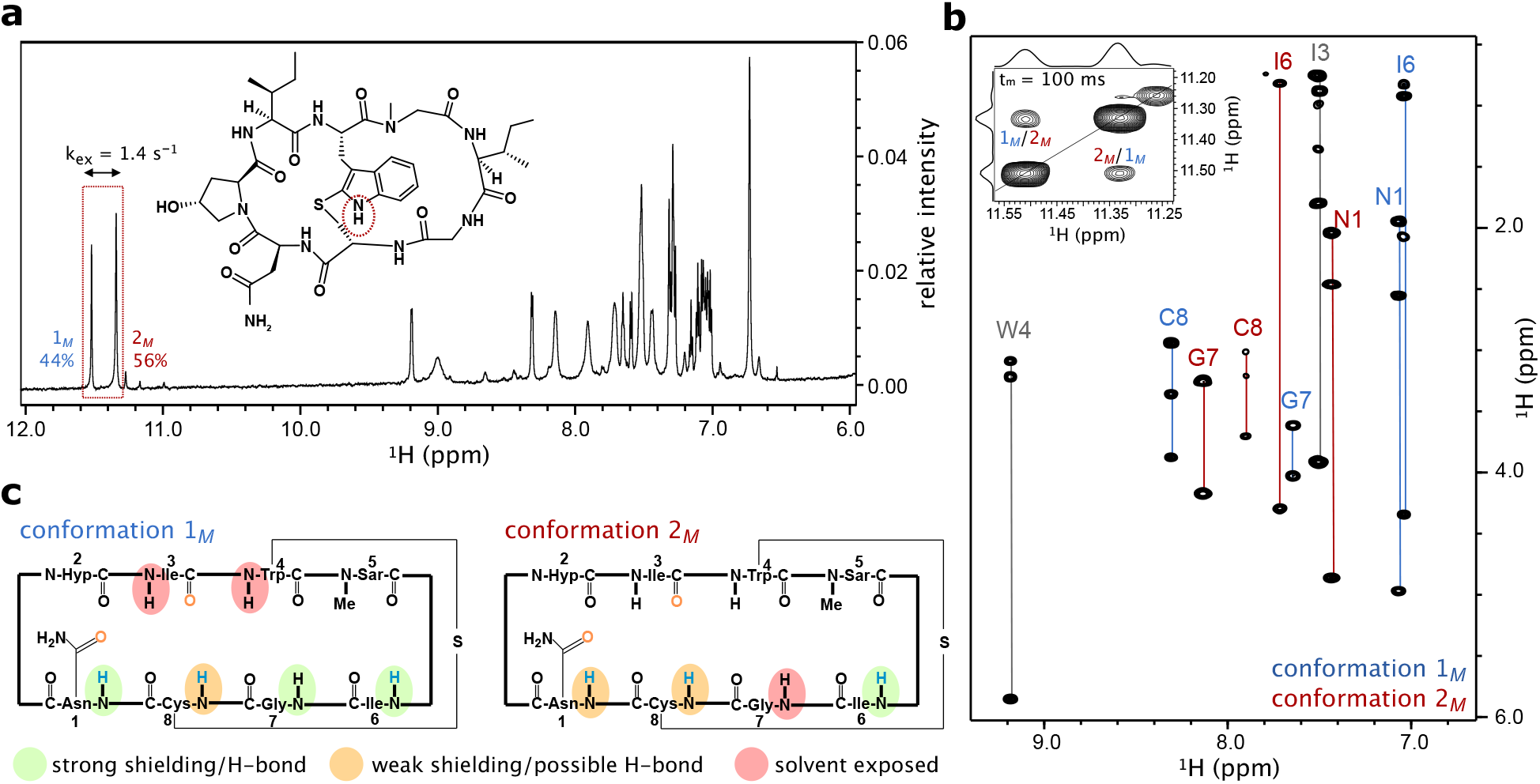
Experimental evidence for the conformational exchange in *M*_ansa_-Gly5Saramanullin. a) Amide region of the ^1^H-NMR spectrum of Gly5Sar-amanullin. Two signals are visible for the indole amine proton (red box), suggesting two conformations of the peptide. The exchange rate *k*_ex_ = 1.4 s^−1^ was measured by EXSY NMR. The rate constants were 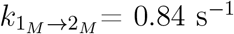 and 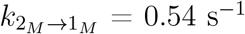. b) The ^1^H-^1^H-TOCSY NMR spectrum of *M*_ansa_-Gly5Sar-amanullin shows two sets of amide correlation signals for Asn1, Ile6, Gly7 and Cys8. The signals were assigned to either conformation 1_*M*_ (blue labels) or conformation 2_*M*_ (red labels). The inset shows the diagonal peaks of the two indole (-NH) proton signals with chemical exchange cross peaks in the TOCSY spectrum. c) Strongly shielded amides with larger temperature correlation coefficients (Δ*δ*HN/Δ*T* > -3.0 ppbK^−1^) suggesting H-bonding are highlighted in green. Solvent exposed amides with small temperature correlation coefficients (Δ*δ*HN/Δ*T* < -4.6 ppbK^−1^) are highlighted in red and amides with weak shielding/H-bonding are shaded in orange.

The ^1^H-^1^H-TOCSY spectrum, two sets of amide correlation signals were observed for Asn1, and the amino acids in the “eastern” ring of the bicycle (Ile6, Gly7 and Cys8). Additionally, we observe an exchange peak between the two indole (-NH) proton peaks (Figure 2.b). The two dominant peaks exchanged with a rate of *k*_ex_ = 1.381 s^−1^ in the ^1^H-^1^H-EXSY spectrum at 298 K.The exchange rate is defined as 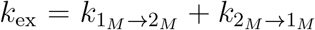, where 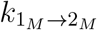 is the rate from 1_*M*_ to 2_*M*_, and 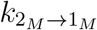 is the rate for the reverse reaction. We measured 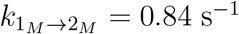 and 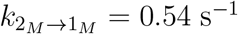 at 298 K. From these rates, we can estimate the populations to be 39% for 1_*M*_ and 61% for 2_*M*_, which is in excellent agreement with the estimates from the 1D-spectrum. On the other hand, a conformational transition that occurs on a timescale of seconds is unusual for a cyclic peptide, which (at least in MD simulations) have conformational equilibration times in the range of 100 ns to several microseconds^30–33^

For both populations, we extracted NOE distances, and we determined the shielding of the amide protons by a variable-temperature NMR experiment (Figure 2.c). The NOE distance set for 1_*M*_ contains 67 upper distance bounds, that of 2_*M*_ contains 41 upper distance bounds. Of these, 11 distances appear in both sets. For 1_*M*_, we could additionally measure the 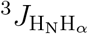-coupling constants which was not possible for 2_*M*_ due to the larger line widths of the amide proton signals. (see Supplementary Table 3).

### Characterisation of *M*_ansa_-Gly5Sar-amanullin

Next, we set out to identify the three-dimensional structure of 1_*M*_ and 2_*M*_ using classical molecular-dynamics (MD) simulations. The exchange rate of *k*_ex_ = 1.381 s^−1^ indicates that, at room temperature, the conformational dynamics is slow compared to the timescales that are accessible by classical MD simulations. The first challenge therefore is to explore the conformational space. Starting form two different structures, we simulated *M*_ansa_-Gly5Saramanullin at 300 K, 400 K, 500 K and 700 K for 24 *μ*s, respectively. The molecule was solvated in dimethyl sulfoxide (DMSO), which was modelled explicitly. The simulations at 400 K and higher temperatures do not represent faithful representations of the molecule at high temperatures but solely serve to explore the conformational space. (See Supplementary Information section 3 for computational details.)

#### Cis-trans isomerism of non-proline residues

At 500 K, the typical, sterically-allowed regions of the Ramachandran plot of each of the residues are explored, and we observe transitions between the two starting conformations.

Additionally, we observe cis-trans isomerisation in several *ω*-torsion angles (Figure 3), specifically *ω*_4_ (between Trp4 and Sar5), *ω*_5_ (between Sar5 and Ile6), *ω*_7_ (between Gly7 and Cys8), and *ω*_8_ (between Cys8 and Asn1). Following the IUPAC definition,^45^ *ω*_*i*_ is the torsion angle defined by C_*α*,i_− carbonyl-C_*i*_− N_*i*+1_ − C_*α,i*+1_. The torsion angle *ω*_1_ (between Ans1 and Hyp2) samples a very broad distribution in the cis-configuration, but does not fully transition to the trans-configuration. The flexibility of *ω*_1_ at high temperatures, however, is not surprising, because Hyp2 is a hydroxylated proline-variant and the peptide group prior to a proline-residue has a relatively small barrier for this isomerisation compared to regular peptide group.^46,47^ Typical isomerisation barriers for a peptide group prior to a prolineresidue are about 90 kJ*·*mol^−1^ Apart from *ω*_1_, the four torsion angles which show cis-trans isomerisation stand out, because they are close to the tryptathionine bridge.

**Figure 3:**
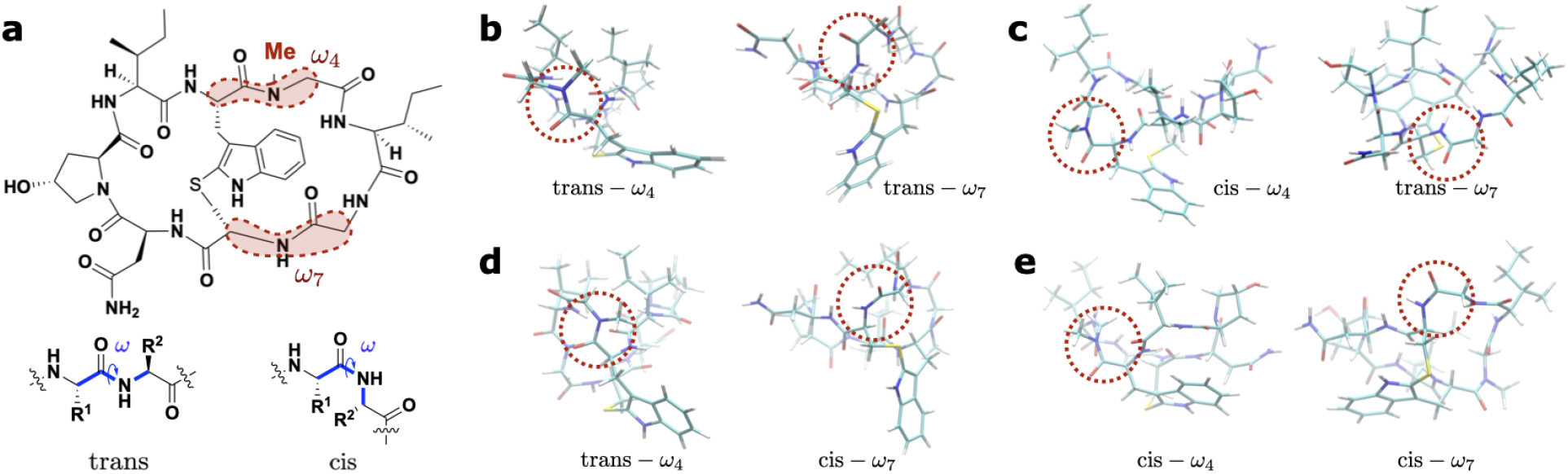
cis-trans isomerism in *M*_ansa_-Gly5Sar-amanullin (a) resulting in four different configurations (b-d). a) chemical structure of Gly5Sar-amanullin with *ω*_4_ and *ω*_7_ highlighted. Below, the general ‘cis’ and ‘trans’ configuration of *ω* are shown for an arbitrary peptide group. *ω* highlighted in blue, is defined according to IUPAC: C_*α*,i−_ carbonyl-C_*i*_-N_*i*+1_-C_*α,i*+1_.^45^ b)-d) For each configuration, the determining (*ω*_4_,*ω*_7_)-torsion angles are shown in the structures (red circles): b) (trans-*ω*_4_, trans-*ω*_7_); c) (cis-*ω*_4_, trans-*ω*_7_); d) (trans-*ω*_4_, cis-*ω*_7_); e) (cis-*ω*_4_, cis-*ω*_7_). Each structure is coloured according to atom type: *cyan:* carbon; *white:* hydrogen; *red:* oxygen; *blue:* nitrogen; *yellow:* sulphur.

We simulated *M*_ansa_-Gly5Sar-amanullin at 700 K to test whether *ω*_2_, *ω*_3_, and *ω*_6_ could also exhibit cis-trans isomerization. This was not the case. Even at this high temperature, the three torsion angles remained in the trans-configuration.

When cooling the ensemble from 500 K to 400 K, we keep observing cis-trans transitions in *ω*_4_ (between Trp4 and Sar5) and *ω*_7_ (between Gly7 and Cys8). *ω*_5_ (between Sar5 and Ile6) and *ω*_8_ (between Cys8 and Asn1) do not show any transitions and only sample the trans-configuration. Cooling the ensemble further to 300 K, we keep observing occasional cis-trans transitions in *ω*_4_. For *ω*_7_, both cisand trans-configurations are present in the ensemble, but in different trajectories. We do not observe any transitions between these two configurations at room temperature during our simulations. (See Supplementary Figures 6-8 for the Ramachandran plots and the distributions of the *ω*-torsion angles at all simulated temperatures as well as the time series of *ω*_4_ at 300 K.)

In the simulated ensemble at 300 K, four different configurations are present (Figure 3): (trans-*ω*_4_, trans-*ω*_7_), (trans-*ω*_4_, cis-*ω*_7_), (cis-*ω*_4_, trans-*ω*_7_), and (cis-*ω*_4_, cis-*ω*_7_). (trans-*ω*_4_, trans-*ω*_7_) is the expected conformation and occurs in the majority of the 24 trajectories. (cis*ω*_4_, trans-*ω*_7_) occurs in two trajectories, and (trans-*ω*_4_, cis-*ω*_7_) and (cis-*ω*_4_, cis-*ω*_7_) each occur in one trajectory (see Supplementary Table 5). Apart from short excursion from (trans-*ω*_4_, trans-*ω*_7_) to (cis-*ω*_4_, trans-*ω*_7_), we don’t observe transitions between the four configurations. Note that we do not observe any cis-trans isomerisation for *ω*_1_, i.e. in the peptide bond which precedes the proline-variant Hyp2.

The chemical shifts of the C_*β*_ proton in Hyp2 of *M*_ansa_-Gly5Sar-amanullin are similar to those in the *M* -asamer of amanullin.^29^ It thus seem likely that *ω*_1_ is in the cis-configuration, which agrees with our MD results. Unfortunately, the configuration of *ω*_4_ and *ω*_7_ cannot be unambiguously assigned from our NMR data. However, cis-trans isomerisation in peptides with N-methylated amino acids and particular in peptides with sarcosine have been reported previously.^48^ Likewise cis-trans isomerisation in strained cyclic peptides without N-methylated amino acids are known.^49^ The exchange rates for the cis-trans isomerisation reported in Ref. 49 are on the order of 1 s^−1^ and are thus in the same range as the exchange rate between 1_*M*_ and 2_*M*_. Overall, it seems possible that 1_*M*_ and 2_*M*_ correspond to two different configurations of the peptide backbone *M*_ansa_-Gly5Sar-amanullin rather than to two different conformations within the (trans-*ω*_4_, trans-*ω*_7_)-ensemble.

#### Comparison of MD structure and NMR data for the *M* -ansamer

We compared the four (*ω*_4_, *ω*_7_)-configurations to our NMR data. The (trans-*ω*_4_, trans*ω*_7_)-MD ensemble shows an almost perfect fit to the NOE upper distance bounds for 1_*M*_ (Figure 4.d). We can only identify two small violations for the 67 NOE distances. The corresponding distances are shown as pink lines in Figure 4.c. By contrast, the other three configurations show considerably more violations of the NOE upper distance bounds for 1_*M*_, in particular for main chain-main chain and the main chain-side chain distances (Supplementary Figure 9).

**Figure 4:**
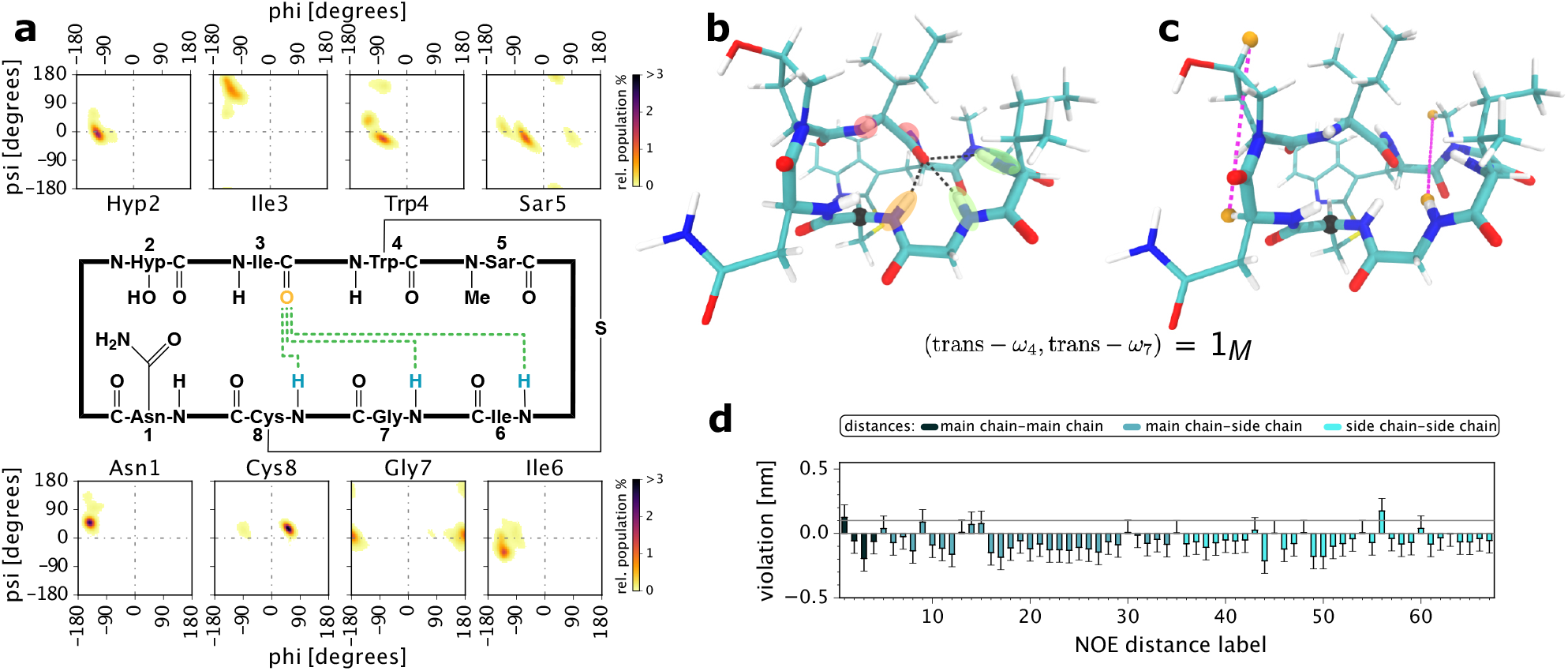
Conformation *M*_ansa_-Gly5Sar-amanullin extracted from the (trans-*ω*_4_, trans-*ω*_7_)- MD ensemble at 300 K. (a) Hydrogen bond pattern and Ramachandran plots. *green dashed line:* hydrogen bonds with ≥ 20% population in the MD ensemble. The colourmap in the Ramachandran plots shows the total probability density within the (trans-*ω*_4_, trans-*ω*_7_)-ensemble. (b,c) MD structure for (trans-*ω*_4_, trans-*ω*_7_)-Gly5Sar-amanullin showing the highest-populated hydrogen bonds (b, black dashed line) or the NOE violations (d, purple dashed lines). The structures are coloured according to atom type. *cyan:* carbon; *white:* hydrogen; *red:* oxygen; *blue:* nitrogen; *yellow:* sulphur. The C_*α*_-atom of Cys8 is highlighted as black sphere. In b), the amides of Ile3, Trp4, Ile6, Gly7 and Cys8 are additionally coloured according to their NMR shielding (Figure 2.c). (d) Violations of the NOE upper distance bounds from the experimental structure 1_*M*_ shown the for the (trans-*ω*_4_, trans-*ω*_7_)-MD ensemble at 300 K. For each NOE distance, the violation is shown as average (bar) and standard deviation (errorbar) over the (trans-*ω*_4_, trans-*ω*_7_)-MD ensemble.

Furthermore, the (trans-*ω*_4_, trans-*ω*_7_)-configuration is stabilised by hydrogen bonds from the amide protons of Ile6, Gly7 and Cys8 to the carbonyl oxygen of Ile3 (Figure 4.a). These hydrogen bonds agree well with the solvent exposure determined from variable-temperature NMR experiments (Figure 2.c and Figure 4.b). The hydrogen-bond pattern of (trans-*ω*_4_, cis-*ω*_7_)-configuration shows a similar agreement with the variable-temperature NMR experiments, but the configurations in which *ω*_4_ is in the cis-configuration are stabilized by hydrogen bond patterns that are not consistent with these data (Supplementary Figures 10-11).

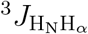-coupling constants depend on the H^*N*^ -C-C-H^*α*^ torsion angle, which related *ϕ*-backbone torsion angle by a shift of about 60°(depending on the bond angles at the two carbon atoms). These coupling constants can thus be used to gauge the conformation of the the H^*N*^ -C-C-H^*α*^ torsion angle and the closely related *ϕ*-backbone angles. (*ϕ*_*i*_ is defined by the backbone atoms C_*i*−1_, N_*i*_, C_*i*[*α*]_, C_*i*_, see Ref.^45^). However, the comparison between simulated and experimental 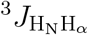-coupling is notoriously difficult, because the Karplus curve which links the *ϕ*-torsion angles to the 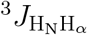 constant is highly-nonlinear and sensitive to the parametrization of the curve. Consequently, inaccuracies of in the potential energy of the simulations can lead to shifts in the predicted 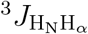 by more than 1 Hz.^50,51^ The comparison of the four configurations to the 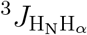-coupling constants for 1_*M*_ confirms that the cis-*ω*_4_-configurations are not consistent with the NMR data of 1_*M*_. Both trans-*ω*_4_-configurations agree equally well with the 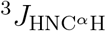-coupling constants for 1_*M*_. We tested different parametrisations of the Karplus curve^52,53^ and accounted for the sensitivity of the estimated 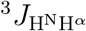 -coupling constants with respect to inaccuracies of the potential energy function by shifting the values of H^*N*^ -C-C-H^*α*^-torsion angle by +10° and -10° (Supplementary Figure 12). Overall, (trans-*ω*_4_, trans-*ω*_7_)-configuration agrees best with the NMR data for 1_*M*_, and we assign this configuration (Figure 4) to the corresponding 1_*M*_ signal in the NMR data.

For 2_*M*_, we have less NMR data, and the comparison to the four configurations unfortunately remains inconclusive. All four configurations show violations of the NOE upper distance bounds for 2_*M*_ (Supplementary Figure 9). However, the trans-*ω*_7_-configurations show fewer violations than the cis-*ω*_7_-configurations. Specifically, they do not have any violations in the critical main-chain-main-chain distances. However, none of the four configurations showed a hydrogen bond pattern that is consistent with the shielding derived of the amide protons in 2_*M*_ as measured in the variable-temperature experiments (Supplementary Figure 11). Overall, 2_*M*_ seems to be consistent with a trans-*ω*_7_-configuration, but a definite assignment is not possible.

Thus, from experimental data it seems likely that *ω*_7_ is in the trans-configuration in 1_*M*_ as well as in 2_*M*_. Since we are confident that *ω*_4_ is in the trans-configuration in 1_*M*_, there are two possible structural explanations for the exchange between 1_*M*_ and 2_*M*_. The exchange can either correspond to a cis-trans isomerisation in *ω*_4_ or to a conformational transition within the (trans-*ω*_4_, cis-*ω*_7_)-configuration.

#### Conformational ensemble of the (trans,trans)-configuration

We analyzed the ensemble of the (trans,trans)-configuration of *M*_ansa_-Gly5Sar-amanullin at 300 K in more detail (Supplementary Figure 13). To identify conformations within this ensemble, we performed a dimensionality reduction based on the conformational dynamics of all backbone torsion angles (time-independent component analysis^54,55^) and clustered in the reduced space using the density-based cluster algorithm common-nearest-neighbour clustering (CommonNN).^56–59^ We obtained 5 clusters (c1-c5, Figure 5.a). Figure 5.b shows the distribution of the ensemble in the reduced space of four time-independent components (TICs, see also Supplementary Figure 14). Figure 5.c shows how the five clusters are located within this space.

**Figure 5:**
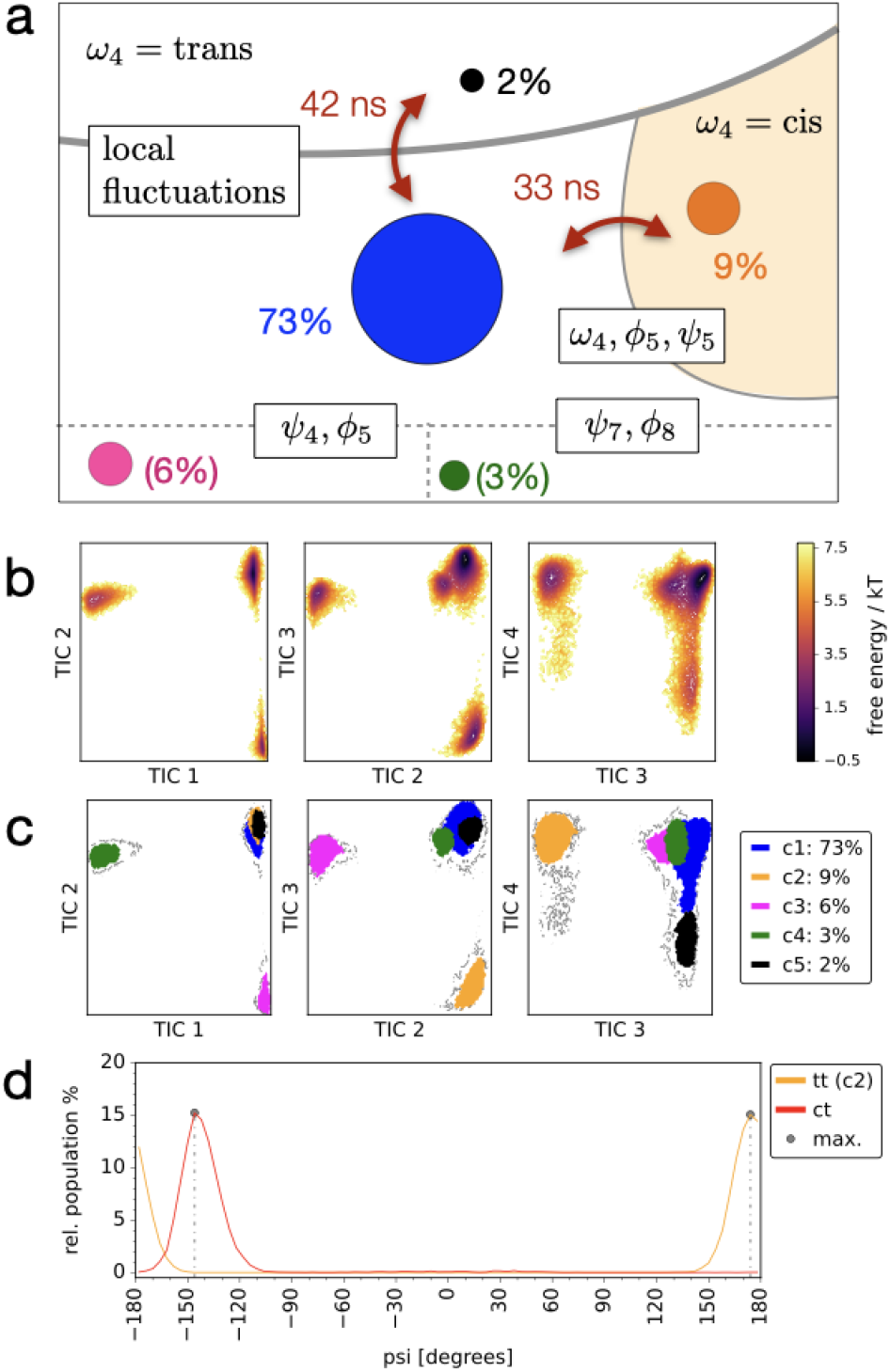
Conformational dynamics of the (trans-*ω*_4_, trans-*ω*_7_)-configuration **a**: kinetic model derived from a core-set Markov model. Clusters c1 to c5 are represented as coloured circles (blue: c1, orange: c2, pink: c3, green: c4, black: c5). Cluster populations are given in % next to the circles. Gray line represent free-energy barriers for the conformational exchange between clusters or groups of clusters. The associated equilibration timescales are shown in red next to the double arrors. White boxes show the torsion angles involved in a particular conformational exchange. **b**: dimensionality reduction by time-independent component analysis: free energy as a function of the first four time-independent components; **c**: CommonNN-clustering in the space of the time-independent components; **d**: distribution of *Ψ*_5_ in the cluster c2 and in the (cis-*ω*_4_, trans-*ω*_7_)-configuration

We determined the timescale of the conformational exchange between these five clusters using a core-set Markov model^58–61^ (Figure 5.a). We find that three clusters (c1, c2 and c5) are well-connected on the timescale of the MD simulation, whereas we do not observe sufficiently many transitions to clusters c3 and c4 to include them in then in the Markov model.

The most populated cluster is c1 (population 73%; blue circle in Figure 5.a) and its structure is shown Figure 4.b. Cluster c5 (population 2%; black circle in Figure 5.a) represents local fluctuations around this structure that do not involve any major transitions in the backbone torsion angles. These local fluctuations equilibrate on a timescale of about 42 ns (Supplementary Figure 15). In both c1 and c5, the *ω*_4_-torsion angles remain in the trans configuration.

By contrast, cluster c2 (population 9%; orange circle in Figure 5.a) represents short excursions into the cis-configuration of *ω*_4_, i.e. the *ω*-torsion angle between Trp4 and Sar5. This transition is coupled to a transition in the *ϕ*and *Ψ*-backbone torsion angle of Sar5. However, the resulting (cis-*ω*_4_,trans-*ω*_7_)-configurations are only stable on a timescale of 33 ns. This timescale is at variance with the observation that the (cis-*ω*_4_,trans-*ω*_7_)-configurations is stable over at least 1 *μ*s in the trajectories discussed above. It seems likely that the excursion into the cis-configuration of *ω*_4_ in cluster c2 are recrossing events.^62^ This means, the molecule crosses the barrier between cis- and trans-configurations, but does not fully reach free energy minimum of the (cis,trans)-configuration and instead quickly returns to (trans,trans)-configuration. This interpretation is substantiated by the fact that the distribution of *Ψ*_5_ in the (cis,trans)-configuration is slightly shifted with respect to the *Ψ*_5_-distribution in the (trans,trans)-configuration. That is, to fully reach a stable (cis,trans)-configuration, the *Ψ*_5_-torsion angle needs to shift by about 40°from 174° in cluster c2 to -146° in the (cis,trans)-ensemble (Figure 5.d). It should be noted, however, that the torsion angles are periodic, so the shift is effectively 40°.

Cluster c3 (population 6%; pink circle in Figure 5.a) represents a transition from cluster c1 in the *Ψ*-torsion angle of Trp4 and the *ϕ*-torsion angle in Sar5, where the intermediate *ω*_4_ remains in the trans-configuration. Cluster c4 (population 3%; green circle in Figure 5.a) represents a transition from cluster c1 in the *Ψ*-torsion angle of Gly7 and the *ϕ*-torsion angle in Cys8, where the intermediate *ω*_7_ remains in the trans-configuration.

Note though that the transitions between c1 and c3 and c4, respectively, are not sampled sufficiently. Thus, the relative populations are not well-converged and are denoted in brackets. (See Supplementary Figure 16-17 for the distributions of the residue-specifc *ω*-backbone torsion angles as well as the hydrogen bonds in the clusters c1-c5.)

### Characterisation of *P*_ansa_-Gly5Sar-amanullin

We explored the conformational space of *P* -ansamer of Gly5Sar-amanullin in a similar manner as for the *M* -ansamer: we simulated the molecule at high temperatures (up to 800 K) and stepwise cooled the ensemble to 300 K. In contrast to *M*_ansa_-Gly5Sar-amanullin, we do not observe any cis-trans isomerisation in the backbone of the *P* -ansamer of Gly5Saramanullin at 300 K. All *ω*-torsion angles are in trans-configuration, which is line with the previously published crystal structure of *P* -amanullin.^29^ However, when cooling the ensemble to 300 K, we find two different conformations of *P*_ansa_-Gly5Sar-amanullin: 1_*P*_ (Figure 6.a) and 2_*P*_ (Figure 6.b). In total, we have 36 *μ*s MD trajectories of 1_*P*_ at temperatures between 300 K and 340 K, and 50 *μ*s MD trajectories of 2_*P*_ at 300 K, but we do not observe any transitions between the two conformations. That is, the conformational exchange between these two structures is slow in the timescale of the MD simulations. 1_*P*_ and 2_*P*_ differ in the conformation of the backbone torsion angle *ϕ* and *Ψ* of most residues and in the intramolecular hydrogen bond pattern (Supplementary Figure 18). In 1_*P*_, the intramolecular hydrogen bond network mostly affects the left cyclic subunit, and consequently the right cyclic subunit is somewhat flexible, in particular in the backbone of Ile6 and Gly7. By constrast, the hydrogen bond network in 2_*P*_ affects all regions of the peptide ring and this conformer is more rigid than 1_*P*_.

**Figure 6:**
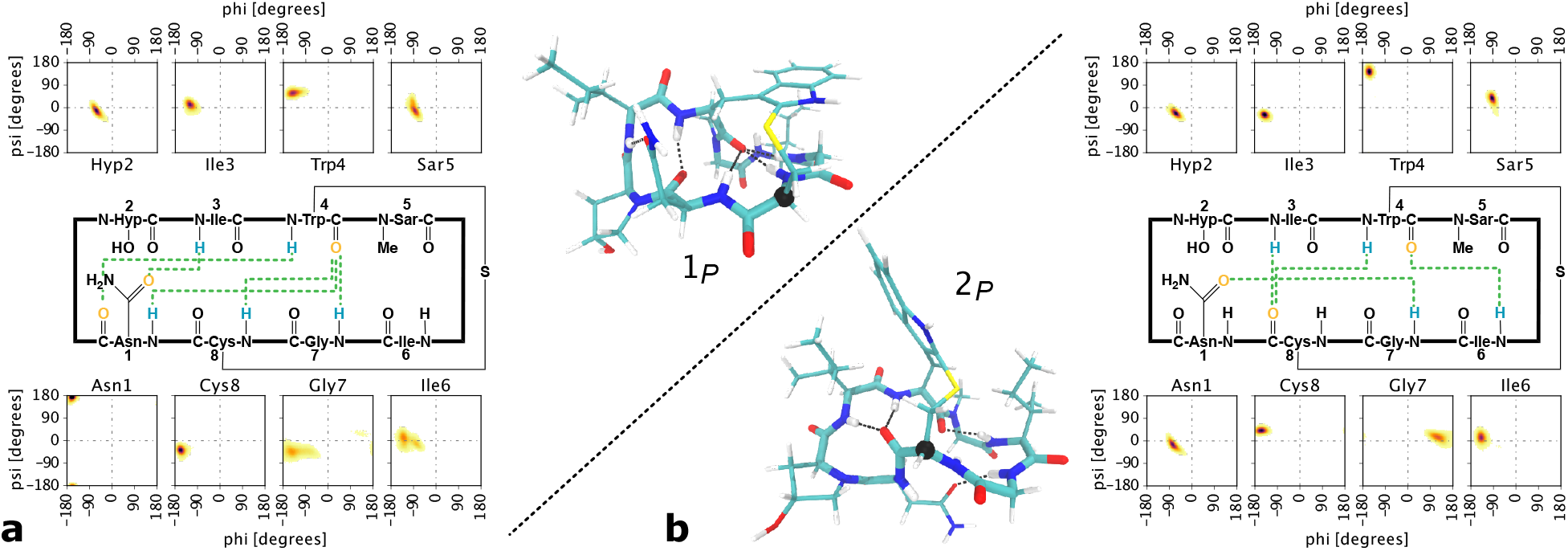
Conformation 1_*P*_ (a) and 2_*P*_ (b) of *P*_ansa_-Gly5Sar-amanullin extracted from MD simulations at 300 K. For each conformation, the hydrogen bond pattern, Ramachandran plots and structure are shown. Hydrogen bonds with a population ≥ 20% within the MD ensemble of the respective conformation are depicted as *green dashed lines* in the scheme and *black dashed lines* in the structure. The structures are coloured according to atom type. *cyan:* carbon; *white:* hydrogen; *red:* oxygen; *blue:* nitrogen; *yellow:* sulphur. The C_*α*_-atom of Cys8 is highlighted as black sphere.

### Comparison between amanullin and Gly5Sar-amanullin

We previously published the crystal structures of both ansamers of amanullin.^29^ Figure 7 compares these structures to the corresponding structures of Gly5Sar-amanullin, which we obtained from our simulations. For Gly5Sar-amanullin, we here only consider the (trans,trans)configuration.

**Figure 7:**
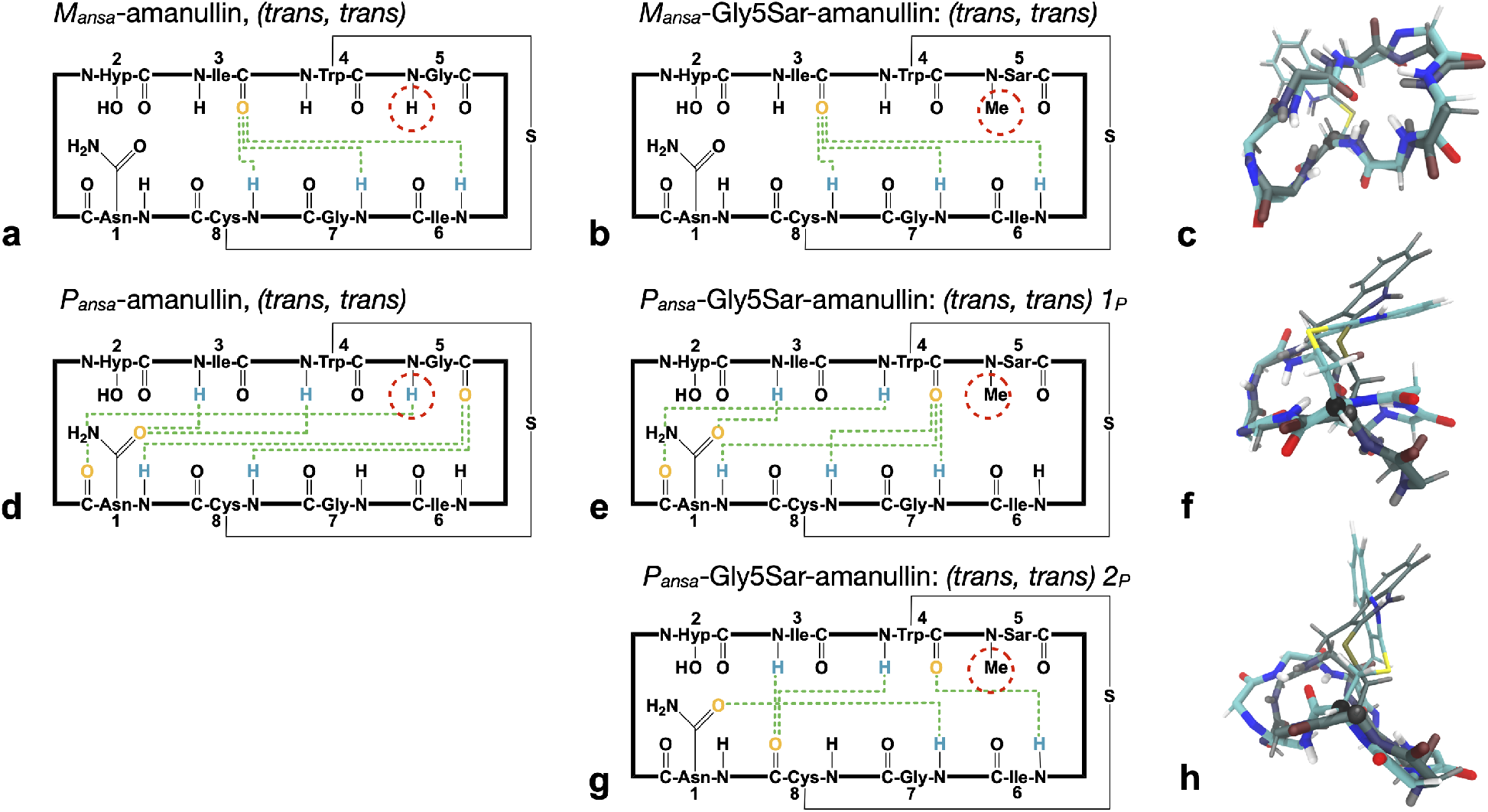
Comparison between the crystal structures for *P*_ansa_-and *M*_ansa_-amanullin (CCDC deposition numbers: 1128063,^41^ 2153904^29^) and the conformations 1_*M*_, 1_*P*_ and 2_*P*_ from our MD simulations on Gly5Sar-amanullin. (a-g) Hydrogen bond schemes for the crystal structures and the structures from MD. Hydrogen bonds are depicted as green dotted lines between the donor atom (blue) and the acceptor atom (orange). (c,f,h) Superposition of the Gly5Sar-amanullin conformations (coloured) 1_*M*_ (c), 1_*P*_ (f) and 2_*P*_ (h) with the respective amanullin crystal structure (grey). The structures are coloured according to atom type: carbon : cyan, hydrogen : white, oxygen : red, nitrogen : blue, sulphur : yellow. All structures have been fitted on the backbone of the residues Gly7, Cys8, Asn1, Hyp2 and Ile3.

The *M* -ansamer of amanullin is stabilized by hydrogen bonds from amide hydrogens of Ile6, Gly7 and Cys8 across the peptide ring to the carbonyl group of Ile3 (Figure 7.a). The amide hydrogen of Gly5 is solvent exposed, and is not involved in stabilizing the structure. Consequently, methylating this position has little effect on the conformation of the molecule. The (trans,trans)-configuration of *M* -Gly5Sar-amanullin exhibits the same hydrogen bond patterns as *M* -amanullin (Figure 7.b), and have the same backbone structure (within the variation due to thermal vibrations)(Figure 7.c).

By contrast, the *P* -ansamer of amanullin is stabilized by a hydrogen-bond network, which includes a hydrogen bond from the amide hydrogen of Gly5 across the peptide ring to the carbonyl oxygen Asn1 (Figure 7.d). This hydrogen bond cannot be formed in *P* -Gly5Saramanullin, and different hydrogen bond patterns arise. Instead of one conformational isomer in *P* -amanullin, we find two major conformations in *P* -Gly5Sar-amanullin. In the conformation 1_*P*_, the hydrogen bond pattern is shifted by one residue compared to *P* -amanullin (Figure 7.e). The Gly5-Ans1 hydrogen bond is replaced by Trp4-Ans1 hydrogen bond. And instead of forming hydrogen bonds to the carbonyl oxygen of Gly5, the amide protons of Asn1 and Cys8 now form hydrogen bonds to Trp4. The conformation 2_*P*_ is stabilized by a hydrogen bond pattern (Figure 7.g) that is completely different from the hydrogen bonds of *P* -amanullin. Consequently, the backbone structure of either conformation of *P* -Gly5Sar-amanullin differs from the conformation of *P* -amanullin (Figures 7.f and 7.h).

To assess possible differences in the physico-chemical properties of the five structures in Figure 7, we calculated their surface accessible area (SASA) and decomposed this area into a hydrophilic and a hydrophobic contribution (Table 1). However, the various structures showed only small differences in these properties.

**Table 1:** Solvent accessible surface area (SASA), hydrophobic and hydrophilic fraction of the SASA

|  |  | SASA [nm<sup>2</sup>] | hydrophilic | hydrophobic |
| --- | --- | --- | --- | --- |
| amanullin |  |  |  |  |
| *M* ansamer | (trans-$\omega_4$,trans-$\omega_7$) | 9.22 | 31 % | 69 % |
| *P* ansamer | (trans-$\omega_4$,trans-$\omega_7$) | 10.05 | 32 % | 68 % |
| Gly5Sar-amanullin |  |  |  |  |
| *M* ansamer | (trans-$\omega_4$,trans-$\omega_7$) | 9.89±0.15 | 37 % | 63 % |
| *P* ansamer | (trans-$\omega_4$,trans-$\omega_7$) $1_P$ | 9.7±0.38 | 32 % | 68 % |
| *P* ansamer | (trans-$\omega_4$,trans-$\omega_7$) $2_P$ | 9.3±0.28 | 34 % | 66 % |

### Conclusion and outlook

We combined NMR experiments and MD simulations to elucidate the influence of N-methylation on the structural ensemble of an amatoxin, specifically we compared Gly5Sar-amanullin and amanullin. Due to their bicyclic structure, amatoxins can form two different ansamers, and the effect of N-methylation is different in the two ansamers.

In the *P* -ansamer of Gly5Sar-amanullin, the methyl group prevented a hydrogen bond that is present in *P*_ansa_-amanullin and two new conformations could arise. The exchange of these two new conformations was slow on the timescale of the MD simulations (microseconds).

By contrast, *M*_ansa_-amanullin does not have a hydrogen-bond to the amide-proton of Gly5, and it is overall very rigid. Thus, one could have expected that methylation of Gly5 has little effect on the structure of this ansamer. Yet, we found two populations in the NMR experiments which exchanged with a rate of 1.38 s^−1^.

The MD-data showed cis-trans isomerization in two *ω*-torsion angles of *M*_ansa_-Gly5Saramanullin: one involved the methylated amide (*ω*_4_), the other was located on the opposite site of the tryptathionine bridge (*ω*_7_). We could assign the (trans-*ω*_4_, trans-*ω*_7_)-configuration to the 1_*M*_ signal in the NMR data. The 2_*M*_ signal could not be assigned unambiguously to an MD-structure. However, we could rule out that 2_*M*_ is a cis-*ω*_7_-configuration. Since the MD simulations of the (trans-*ω*_4_, trans-*ω*_7_)-configuration showed short excursions to the (cis-*ω*_4_, trans-*ω*_7_)-configuration, it seems likely that the exchange between the two NMR signals corresponds to a cis-trans isomerization

Our results show that, even though amatoxins have a relatively rigid scaffold, methylation of the backbone amides can have varied and sizeable effects on their overall structure. Nmethylation can lower the rotational barrier of the corresponding *ω*-torsion angle and can give rise to thermal cis-trans isomerization on the timescale of seconds. And even within a given configuration, the introduction of a methyl-group prohibits certain intramolecular hydrogen bonds and generates new conformations.

## Methods

Computational and experimental methods are reported in the Supplementary Information.

## Acknowledgement

This research has been funded by Deutsche Forschungsgemeinschaft (DFG) through grant RTG 2473 Bioactive Peptides - Project ID 392923329 - Project C01. SFB 1349 Fluorine- Specific Interactions - Project ID 387284271, Project A05. We thank the Zentraleinrichtung für Datenverarbeitung (ZEDAT) of Freie Universität Berlin for computing time. We also thank the NMR Messzentrum at the Institute of Chemistry at TU Berlin for the use of their NMR spectrometers and especially Samantha Voges for assistance with spectra acquisition. Guiyang Yao thanks the Alexander von Humboldt Foundation for a postdoctoral fellowship and the National Natural Science Foundation of China (82204189).

## Supporting Information Available

We provide the protocols (section 1) as well as spectroscopic evidence (section 2.1) for the synthesis of *M*_ansa_-Gly5Sar-amanullin: reagents, solvents, chromatographic conditions, UV- VIS-, ^1^H-NMR- and CD spectra.

We also provide all evidence from NMR experiments that indicates the presence of two long-living conformations for *M*_ansa_-Gly5Sar-amanullin (1_*M*_, 2_*M*_, section 2.2): chemical shifts, amide temperature coefficients, 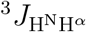-coupling coefficients and inter-proton distances from ^1^H-^1^-NOESY-spectra (NOE distances)

Furthermore, we provide additional material for the characterisation of (1) the MD ensembles of *M*_ansa_- and *P*_ansa_-Gly5Sar-amanullin at different temperatures (2) the sub data sets for the 300 K MD ensemble of the *M* -ansamer incl. the assignment of the trajectories and the clusters from sub data set ‘tt’ of the 300 K MD ensemble of *M*_ansa_-Gly5Sar-amanullin (section 2.3) The characterisation involves backbone torsion angle (*ϕ, Ψ, ω*) distributions, calculated 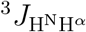-coupling coefficients using different parameter sets for the Karplus equation, hydrogen bond populations and schemes with amides coloured according to expected solvent exposure, analysis of the inter-proton distances from the experimental NOE distances for 1_*M*_ and 2_*M*_ (tables and violation plots) and, for the clustering of the ‘tt’ sub data set: the eigenvalues and distributions for the time-independent components as well as the implied timescales for the constructed cs-MSM.

In sections 3 and 4, all computational and experimental methods used in this study are described and necessary parameters are provided. This involves the setup for the MD simulations of both ansamers of Gly5Sar-amanullin as well as the computational tools for their analysis (section 3). In addition, further details on the HPLC-MS, CD spectroscopy and on the NMR experiments (incl. proton assignments and variable temperature measurements) are given (section 4).

