## Supplementary Information for "The influence of N-methylation on the ansamers of an amatoxin: Gly5Sar-amanullin"

**References** **48**

### Abbreviations

- CH3CN: acetonitrile
- THF: tetrahydrofuran
- DCM: dichloromethane
- DIPEA: *N*,*N*'-diisopropylethylamine
- DMF: *N*,*N*'-dimethylformamide
- TFA: trifluoroacetic acid
- THF: tetrahydrofuran
- HPLC: high-performance liquid chromatography
- HRMS: high resolution mass spectrometry
- HATU: *O*-(7-azabenzotriazol-1-yl)-*N*,*N*,*N*',*N*'-tetramethyluronium -hexafluorophosphate
- 2-CTC: 2-chlorotrityl chloride (resin); TIS, triisopropylsilane
- HFIP: hexafluoroisopropanol
- Fmoc: fluorenylmethoxycarbonyl protecting group.

The amino acid three letter code was used for the proteinogenic amino acids according to IUPAC standards. If not otherwise stated, L-amino acids were used. Amino acid abbreviations: Hyp, *trans*-4-hydroxy-L-proline.

### 1 Synthesis

#### 1.1 Reagents, Solvents and Chromatographic Conditions

Commercially available reagents (Sigma-Aldrich, Iris Biotech GmbH, ABCR, TCI, VWR) and solvents (Fisher Scientific-Acros) were used without further purification.

#### 1.2 General protocol for solid-phase peptide synthesis (SPPS)

**Method A) Removal of the Fmoc group.** A solution of 20% piperidine in DMF (5 mL) was added to the resin (1 g; loading 0.10-0.50 mmol/g) and the resulting suspension was shaken for 10 min. Then, the solution was removed from the resin. Again, a solution of 20% piperidine in DMF (5 mL) was added to the resin and the resulting suspension was shaken for another 10 min. The solution was drained and the resin was washed with DMF (6 x 5 mL).

**Method B) Amino acid coupling.** Amino acid (4.0 eq) and TBTU (4.0 eq) were dissolved in dry DMF (5 mL). DIPEA (12 eq) was added dropwise to the DMF solution. After activating for 1 min, the resulting solution was added to the Fmoc-deprotected resin (1 g; loading 0.10-0.50 mmol/g). The mixture was shaken until the coupling reaction was completed. Then, the solution was drained and the resin was rinsed with DMF (4 x 5 mL).

#### 1.3 Synthesis of monocyclic peptide 1

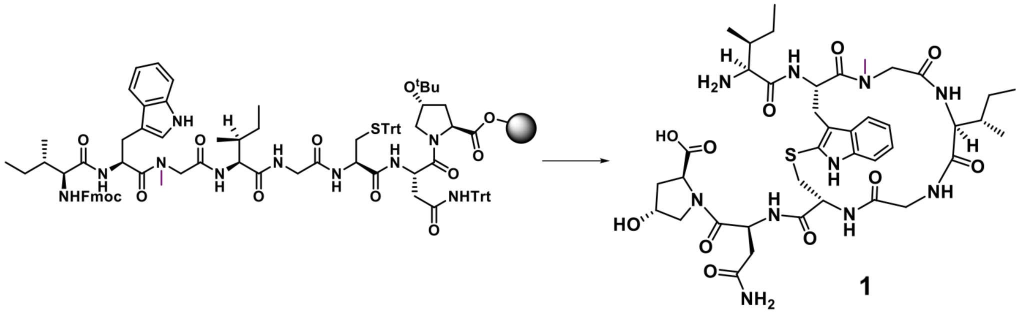

**Supplementary Figure 1.** Synthesis of monocyclic octapeptide H2N-Ile-(*cyclo-tryptathionine*)[Trp-Sar-Ile-Gly-Cys]-Asn-Hyp-OH (**1**).

2-CTC resin (1 g, 0.98 mmol/g) was pre-swollen for 20 min in DCM in a manual solid phase peptide synthesis vessel (10 mL). After the solvent was drained, the first amino acid Fmoc-Hyp(O$^t$Bu)-OH (0.3 mmol) and DIPEA (0.26 mL, 1.5 mmol) in DCM (5 mL) were added to the resin. The mixture was agitated for 2 h before the solvent was drained. The resin was rinsed with DMF (4 x 3 mL). Then, a mixture of MeOH/DIPEA/DCM (1:1:8) was added to cap the remaining 2-chlorotrityl chloride on the resin. The mixture was agitated for 0.5 h. Then, the solvent was drained and the resin was washed with DMF (4 x 3 mL). The resin loading was determined to be 0.30 mmol/g. The Fmoc-group was removed according to Method A. Fmoc-AA2-OH (4 eq) was coupled to the deprotected resin according to Method B. The Fmoc-group of the resulting resin was removed according to Method A. The following six amino acids (Fmoc-Asn(Trt)-OH, Fmoc-Cys(Trt)-OH, Fmoc-Gly-OH, Fmoc-Ile-OH, Fmoc-Sar-OH, Fmoc-Trp-OH and Fmoc-Ile-OH) were coupled to the deprotected resin according to Method A and B. The tryptathionine formation was carried out on the solid support according $I_2$-mediated thioether formation from Ref. 1. After removal of the Fmoc-group using method A and followed by cleavage from the resin, the monocyclic peptide was obtained following subsequent HPLC purification.

#### 1.4 Synthesis of bicyclic peptide 2

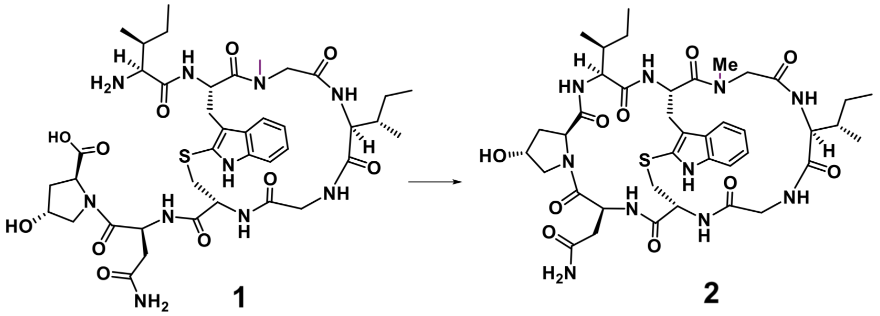

**Supplementary Figure 2.** Synthesis of bicyclic peptide **2** starting from H2N-Ile-(*cyclo-tryptathionine*)[Trp-Sar-Ile-Gly-Cys]-Asn-Hyp-OH (**1**.)

Monocyclic octapeptide **1** (1.0 eq) was dissolved in DMF (1 mM). Then, DIPEA (2.2 eq) and HATU (2.0 eq) was added at 0°C. The reaction mixture was allowed to warm to r.t. for 12 h and concentrated under reduced pressure. The crude product was purified using preparative HPLC to afford bicyclic octapeptide as a white powder.

### 2 Structural charactersiation

#### 2.1 Characterisation of $M_{\text{ansa}}$-Gly5Sar-amanullin and precursor

##### 2.1.1 HRMS and HPLC-MS

H2N-Ile-(*cyclo-tryptathionine*)[Trp-Sar-Ile-Gly-Cys]-Asn-Hyp-OH:

HRMS (ESI): m/z calculated: $C_{40}H_{59}N_{10}O_{11}S^{+}$ $[M+H]^{+}$ 887.4080, found 887.4068.

HPLC-MS: Retention time $R_t = 7.06$ min, $\lambda_{\text{max}}$ 293 nm.

$M_{\text{ansa}}$-Gly5Sar-amanullin:

HRMS (ESI): m/z calculated: $C_{40}H_{57}N_{10}O_{10}S^{+}$, 869.3974, found 869.3970.

HPLC-MS : Retention time $R_t = 10.83$ min, $\lambda_{\text{max}}$ 293 nm.

##### 2.1.2 Spectra

Since UV absorption of tryptathionines is highly distinctive, it has been previously employed to characterize $P_{\text{ansa}}$- and $M_{\text{ansa}}$-amanullin.[2] Interestingly, the UV maximum absorption of peptide **2** is 293 nm, which suggested the $M_{\text{ansa}}$-conformation.

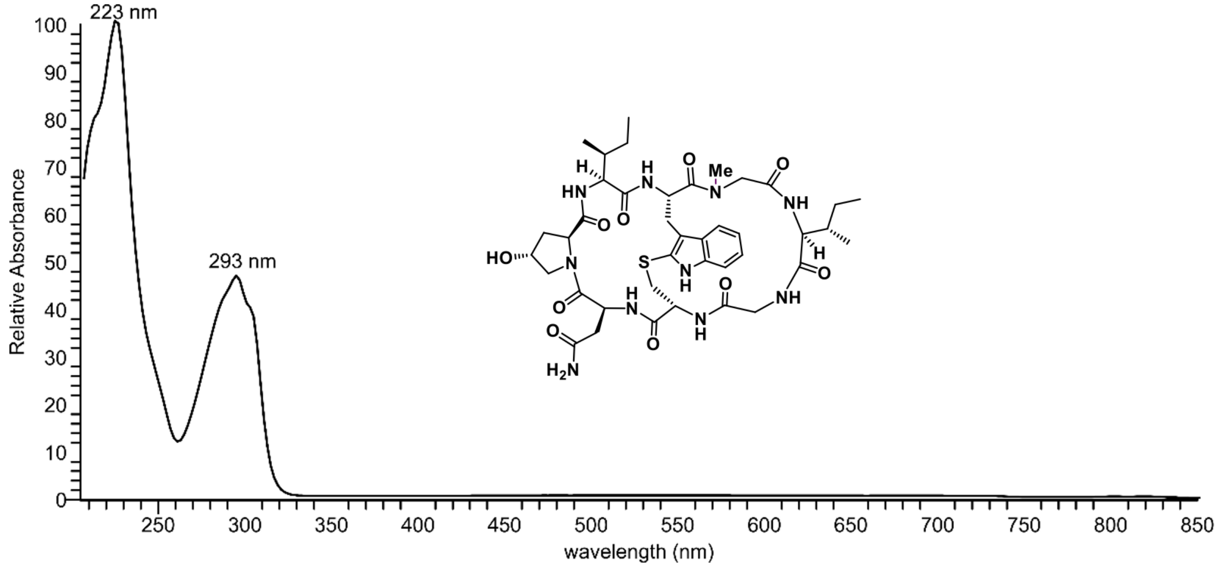

**Supplementary Figure 3.** UV-VIS spectrum of bicyclic peptide **2**.

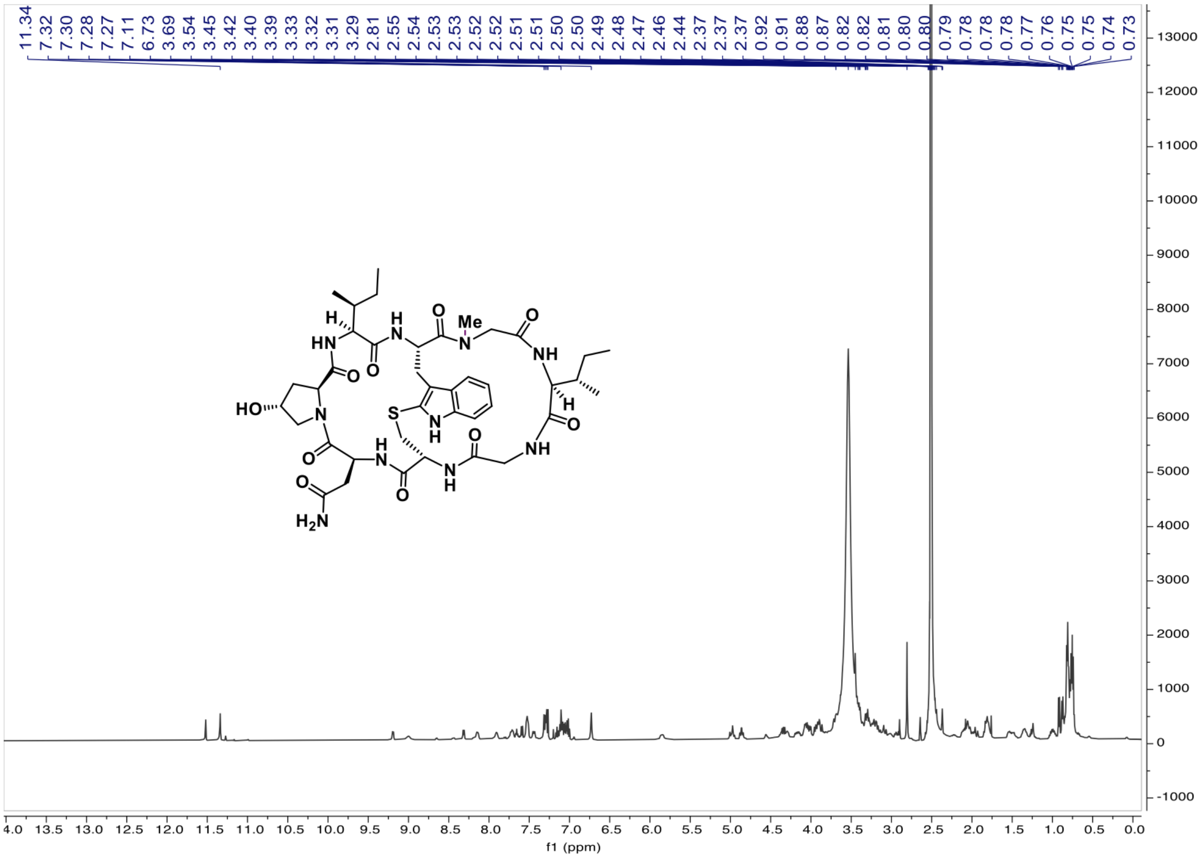

**Supplementary Figure 4.** $^{1}H$ NMR spectrum of bicyclic peptide **2**.

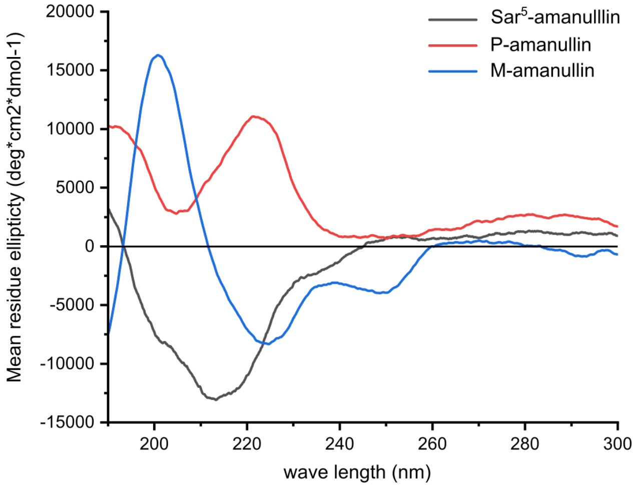

**Supplementary Figure 5.** CD spectra of the $M_{\mathrm{ansa}}$-Gly5Sar-amanullin synthesised in this study (black line, compound **2**), and of the $M_{\mathrm{ansa}}$- and $P_{\mathrm{ansa}}$-isomers for amanullin from our previous study (Ref. 2, red and blue line).

#### 2.2 Data from NMR for the two conformations of $M_{\text{ansa}}$-Gly5Sar-amanullin

**Supplementary Table 1.** Chemical shift assignments in [ppm] of the two conformations of the $M_{\text{ansa}}$-Gly5Sar-amanullin ($1_M$, $2_M$). Methyl groups are indicated by *.

| | | $^1$H Shift (ppm) | | | | | | | | | | | $^{13}$C Shift (ppm) | | | | |
|---|---|---|---|---|---|---|---|---|---|---|---|---|---|---|---|---|---|
| | Residue | H | $H_\alpha$ | $H_\beta$ | $H_\gamma$ | $H_\delta$ | $H_{\epsilon 1}$ | $H_{\epsilon 3}$ | $H_{\zeta 2}$ | $H_{\zeta 3}$ | $H_\eta$ | $H_{n*}$ | $C_\alpha$ | $C_\beta$ | $C_\gamma$ | $C_\delta$ | $C_n$ |
| conformation $1_M$ | Asn1 | 7.07 | 4.97 | 1.95 | | 7.30 | | | | | | | 46.84 | 36.78 | | | |
| | | | | 2.55 | | 6.72 | | | | | | | | | | | |
| | Hyp2 | | 5.11 | 2.11 | 4.06 | 3.41 | | | | | | | | 39.85 | 66.55 | 52.7 | |
| | | | | 1.82 | | 3.30 | | | | | | | | | | | |
| | Ile3 | 7.51 | 3.91 | 1.8 | 0.89* | 0.76* | | | | | | | 56.45 | 34.44 | 23.95 | 9.49* | |
| | | | | | 1.00 | | | | | | | | | | 14.03* | | |
| | | | | | 1.35 | | | | | | | | | | | | |
| | Trp4 | 9.18 | 5.85 | 3.09 | | | 11.52 | 7.59 | 7.31 | 7.05 | 7.16 | | | 22.51 | | | |
| | | | | 3.22 | | | | | | | | | | | | | |
| | Sar5 | | 4.99 | | | | | | | | | 3.45* | 51.29 | | | | 36.34 |
| | | | 3.31 | | | | | | | | | | | | | | |
| | Ile6 | 7.04 | 4.34 | 2.07 | 0.92* | 0.82* | | | | | | | 57.03 | 34.6 | 16.60* | 11.16* | |
| | | | | | 1.50 | | | | | | | | | | | | |
| | Gly7 | 7.64 | 4.03 | | | | | | | | | | 42.64 | | | | |
| | | | 3.61 | | | | | | | | | | | | | | |
| | Cys8 | 8.31 | 2.94 | 3.36 | | | | | | | | | 54.09 | 35 | | | |
| | | | | 3.87 | | | | | | | | | | | | | |
| conformation $2_M$ | Asn1 | 7.43 | 4.83 | 2.01 | | | | | | | | | 46.5 | 36.72 | | | |
| | | | | 2.43 | | | | | | | | | | | | | |
| | Trp4 | | | | | | 11.34 | 7.52 | 7.27 | 7.01 | 7.11 | | | | | | |
| | Ile6 | 7.71 | 3.67 | 2.03 | 1.55 | 0.81* | | | | | | | 56.87 | 33.75 | 23.63 | 10.96* | |
| | | | | | 0.78 | | | | | | | | | | 12.58* | | |
| | | | | | 0.81* | | | | | | | | | | | | |
| | Gly7 | 8.14 | 4.15 | | | | | | | | | | 42.39 | | | | |
| | | | 3.22 | | | | | | | | | | | | | | |
| | Cys8 | 7.9 | 3.22 | 3.01 | | | | | | | | | | | | | |
| | | | | 3.70 | | | | | | | | | | | | | |

**Supplementary Table 2.** Amide temperature coefficients. Shown are the correlation coefficients of the chemical shift changes of the amide protons with increasing temperature. The amide protons with a distinct shift in the second conformation are listed. Colours indicate shielding as follows: green: strong shielding or H-bonding ($> -3.0$ ppbK$^{-1}$), red: solvent exposed ($< -4.6$ ppbK$^{-1}$) and yellow: weak shielding or H-bonding ($< -3.0$ and $> -4.6$ ppbK$^{-1}$).

| | $\Delta\delta$HN / $\Delta T$ (ppbK$^{-1}$) | | | | | |
|---|---|---|---|---|---|---|
| conformation | Asn1 | Ile3 | Trp4 | Ile6 | Gly7 | Cys8 |
| $1_M$ | 1.93 | -9.07 | -4.61 | -0.45 | -2.21 | -3.11 |
| $2_M$ | -4.31 | | | 1.78 | -6.15 | -4.18 |

**Supplementary Table 3.** Experimental $^3J_{\mathrm{H^N-H^\alpha}}$-coupling constants for conformation $1_M$ of Gly5Sar-amanullin. Possible solutions to the Karplus equation (see eq. 3) using the coefficients from Ref. 3 ($A = 6.64$, $B = -1.43$, $C = 1.86$). Note, that for the second conformation ($2_M$), no coupling constants could be obtained due to the line width of amide signals.

| Residue | $^3J_{\mathrm{H^N-H^\alpha}}$ [Hz] | $\phi$ [°] |
|---|---|---|
| Asn1 | 7.03 | -81, -159, 55 |
| Ile3 | 7.75 | -87, -152 |
| Trp4 | 4.82 | -65, -175, 21 |
| Ile6 | 10.11 | -120 |
| Gly7 | -2.21 | – |
| Cys8 | 6.69 | -79, -161, 45 |

**Supplementary Table 4.** Inter-proton distances for the conformations $1_M$ (a) and $2_M$ (b) taken from $^1$H-$^1$H-NOESY spectroscopy. The distances are labelled according to the amino acid sequence: 'Asn1-Hyp2-Ile3-Trp4-Sar5-Ile6-Gly7-Cys8' with the atom names according to standard CYANA nomenclature.[4] Protons that could not be assigned unambiguously are indicated by *. All distances are sorted according to the assignment of the protons to either the main chain ('m', atoms: $H_N$), the side chain ('s'; atoms: else) or (only for Sar5) the main chain II ('x', H of the methyl group replacing $H_N$). All distances are given in [nm]. For a comparison to the distance averages from MD simulations, please refer to Supplementary Table 6.

**(a)** NOE distances for $1_M$.

| ID | type | name | atoms | $R^{\text{NMR}}$ [nm] |
|---|---|---|---|---|
| 1 | x-m | Sar5(Hn*)-Gly7(H) | 106-79 | 0.4 ± 0.16 |
| 2 | m-x | Ile6(H)-Sar5(Hn*) | 74-106 | 0.4 ± 0.16 |
| 3 | m-m | Cys8(H)-Gly7(H) | 82-79 | 0.395 ± 0.158 |
| 4 | m-m | Asn1(H)-Cys8(H) | 86-82 | 0.343 ± 0.137 |
| 5 | s-m | Ile3(Hd1*)-Ile3(H) | 100-96 | 0.38 ± 0.152 |
| 6 | s-m | Ile3(Hg1a)-Ile3(H) | 63-96 | 0.347 ± 0.139 |
| 7 | m-s | Trp4(H)-Ile3(Hg2*) | 64-103 | 0.417 ± 0.167 |
| 8 | m-s | Trp4(H)-Trp4(Ha) | 64-65 | 0.421 ± 0.168 |
| 9 | s-x | Trp4(Ha)-Sar5(Hn*) | 65-106 | 0.318 ± 0.127 |
| 10 | s-m | Trp4(Ha)-Ile6(H) | 65-74 | 0.415 ± 0.166 |
| 11 | s-m | Trp4(Ha)-Cys8(H) | 65-82 | 0.359 ± 0.144 |
| 12 | s-m | Trp4(Hba)-Trp4(H) | 66-64 | 0.408 ± 0.163 |
| 13 | s-m | Trp4(Hbb)-Trp4(H) | 67-64 | 0.351 ± 0.14 |
| 14 | s-m | Trp4(Hbb)-Cys8(H) | 67-82 | 0.405 ± 0.162 |
| 15 | m-s | Ile6(H)-Ile6(Hd1*) | 74-109 | 0.38 ± 0.152 |
| 16 | m-s | Ile6(H)-Ile6(Hb) | 74-76 | 0.429 ± 0.172 |
| 17 | m-s | Ile6(H)-Ile6(Hg1a) | 74-77 | 0.453 ± 0.181 |
| 18 | s-m | Ile6(Ha)-Ile6(H) | 75-74 | 0.406 ± 0.162 |
| 19 | s-m | Ile6(Ha)-Gly7(H) | 75-79 | 0.392 ± 0.157 |
| 20 | s-m | Ile6(Hb)-Gly7(H) | 76-79 | 0.405 ± 0.162 |
| 21 | s-m | Gly7(Haa)-Gly7(H) | 80-79 | 0.36 ± 0.144 |
| 22 | s-m | Gly7(Hab)-Gly7(H) | 81-79 | 0.404 ± 0.162 |
| 23 | m-s | Cys8(H)-Gly7(Haa) | 82-80 | 0.455 ± 0.182 |
| 24 | m-s | Cys8(H)-Gly7(Hab) | 82-81 | 0.45 ± 0.18 |
| 25 | m-s | Cys8(H)-Cys8(Ha) | 82-83 | 0.334 ± 0.133 |
| 26 | m-s | Asn1(H)-Cys8(Ha) | 86-83 | 0.406 ± 0.162 |
| 27 | m-s | Asn1(H)-Asn1(Hba) | 86-88 | 0.428 ± 0.171 |
| 28 | m-s | Asn1(H)-Asn1(Hbb) | 86-89 | 0.376 ± 0.15 |
| 29 | s-m | Asn1(Ha)-Asn1(H) | 87-86 | 0.381 ± 0.152 |
| 30 | s-m | Hyp2(Hg)-Ile3(H) | 93-96 | 0.375 ± 0.15 |
| 31 | m-s | Ile3(H)-Ile3(Hb) | 96-98 | 0.298 ± 0.119 |
| 32 | s-m | Ile3(Ha)-Trp4(H) | 97-64 | 0.295 ± 0.118 |
| 33 | s-m | Ile3(Ha)-Ile3(H) | 97-96 | 0.341 ± 0.136 |

**(a)** (continued) NOE distances for $1_M$.

| ID | type | name | atoms | $R^{\text{NMR}}$ [nm] |
|---|---|---|---|---|
| 34 | s-m | Ile3(Hg1b)-Ile3(H) | 99-96 | 0.326 ± 0.131 |
| 35 | s-s | Ile3(Hd1*)-Ile3(Hb) | 100-98 | 0.286 ± 0.114 |
| 36 | s-s | Ile3(Hg2*)-Ile3(Hb) | 103-98 | 0.334 ± 0.133 |
| 37 | s-s | Ile3(Hg1a)-Ile3(Hd1*) | 63-100 | 0.319 ± 0.128 |
| 38 | s-s | Ile3(Hg1a)-Ile3(Hb) | 63-98 | 0.356 ± 0.142 |
| 39 | s-s | Ile3(Hg1a)-Ile3(Hg1b) | 63-99 | 0.247 ± 0.099 |
| 40 | s-s | Trp4(Hba)-Trp4(Ha) | 66-65 | 0.351 ± 0.14 |
| 41 | s-s | Trp4(Ha)-Cys8(Ha) | 65-83 | 0.348 ± 0.139 |
| 42 | s-s | Trp4(Hba)-Trp4(Ha) | 66-65 | 0.351 ± 0.14 |
| 43 | s-s | Trp4(He1)-Cys8(Hba) | 68-84 | 0.351 ± 0.14 |
| 44 | s-s | Trp4(He1)-Cys8(Hbb) | 68-85 | 0.435 ± 0.174 |
| 45 | s-s | Ile6(Ha)-Ile6(Hd1*) | 75-109 | 0.324 ± 0.129 |
| 46 | s-s | Ile6(Ha)-Ile6(Hg2*) | 75-112 | 0.403 ± 0.161 |
| 47 | s-s | Ile6(Ha)-Ile6(Hb) | 75-76 | 0.34 ± 0.136 |
| 48 | s-s | Ile6(Hb)-Ile6(Hd1*) | 76-109 | 0.289 ± 0.115 |
| 49 | s-s | Ile6(Hb)-Ile6(Hg2*) | 76-112 | 0.432 ± 0.173 |
| 50 | s-s | Ile6(Hb)-Ile6(Hg2*) | 76-114 | 0.432 ± 0.173 |
| 51 | s-s | Gly7(Hab)-Gly7(Haa) | 81-80 | 0.274 ± 0.11 |
| 52 | s-s | Asn1(Ha)-Asn1(Hba) | 87-88 | 0.376 ± 0.15 |
| 53 | s-s | Hyp2(Ha)-Hyp2(Hba) | 90-95 | 0.427 ± 0.171 |
| 54 | s-s | Hyp2(Hg)-Ile3(Hd1*) | 93-100 | 0.383 ± 0.153 |
| 55 | s-s | Hyp2(Hg)-Ile3(Hg1a) | 93-63 | 0.41 ± 0.164 |
| 56 | s-s | Hyp2(Hg)-Asn1(Ha) | 93-87 | 0.382 ± 0.153 |
| 57 | s-s | Hyp2(Hg)-Hyp2(Hbb) | 93-94 | 0.317 ± 0.127 |
| 58 | s-s | Hyp2(Hg)-Hyp2(Hba) | 93-95 | 0.31 ± 0.124 |
| 59 | s-s | Hyp2(Hg)-Ile3(Hg1b) | 93-99 | 0.381 ± 0.152 |
| 60 | s-s | Ile3(Ha)-Ile3(Hd1*) | 97-100 | 0.31 ± 0.124 |
| 61 | s-s | Ile3(Ha)-Ile3(Hg2*) | 97-103 | 0.356 ± 0.142 |
| 62 | s-s | Ile3(Ha)-Ile3(Hg1a) | 97-63 | 0.363 ± 0.145 |
| 63 | s-s | Ile3(Ha)-Trp4(Ha) | 97-65 | 0.422 ± 0.169 |
| 64 | s-s | Ile3(Ha)-Ile3(Hb) | 97-98 | 0.319 ± 0.128 |
| 65 | s-s | Ile3(Ha)-Ile3(Hg1b) | 97-99 | 0.384 ± 0.154 |
| 66 | s-s | Ile3(Hg1b)-Ile3(Hd1*) | 99-100 | 0.295 ± 0.118 |
| 67 | s-s | Ile3(Hg1b)-Ile3(Hb) | 99-98 | 0.332 ± 0.133 |

#### Supplementary Table 4. (continued)

##### (b) NOE distances for $2_M$.

| ID | type | name | atoms | $R^{\mathrm{NMR}}$ [nm] |
|---|---|---|---|---|
| 1 | m-m | Gly7(H)-Cys8(H) | 79-82 | 0.308 ± 0.123 |
| 2 | m-m | Ile6(H)-Gly7(H) | 74-79 | 0.334 ± 0.134 |
| 3 | m-m | Asn1(H)-Cys8(H) | 86-82 | 0.331 ± 0.132 |
| 4 | m-m | Ile6(H)-Cys8(H) | 74-82 | 0.356 ± 0.142 |
| 5 | s-m | Cys8(Hba)-Cys8(H) | 84-82 | 0.369 ± 0.148 |
| 6 | s-m | Cys8(Ha)-Cys8(H) | 83-82 | 0.311 ± 0.124 |
| 7 | s-m | Ile6(Ha)-Ile6(H) | 75-74 | 0.359 ± 0.144 |
| 8 | s-m | Ile6(Ha)-Gly7(H) | 75-79 | 0.333 ± 0.133 |
| 9 | s-m | Asn1(Hbb)-Asn1(H) | 89-86 | 0.363 ± 0.145 |
| 10 | s-m | Ile6(Hg1b)-Ile6(H) | 78-74 | 0.351 ± 0.14 |
| 11 | s-m | Ile6(Hb)-Gly7(H) | 76-79 | 0.416 ± 0.166 |
| 12 | s-m | Ile6(Hb)-Ile6(H) | 76-74 | 0.381 ± 0.152 |
| 13 | s-m | Asn1(Ha)-Asn1(H) | 87-86 | 0.363 ± 0.145 |
| 14 | s-m | Trp4(Ha)-Cys8(H) | 65-82 | 0.43 ± 0.172 |
| 15 | m-s | Cys8(H)-Gly7(Hab) | 82-81 | 0.371 ± 0.148 |
| 16 | m-s | Gly7(H)-Gly7(Hab) | 79-81 | 0.335 ± 0.134 |
| 17 | m-s | Gly7(H)-Gly7(Haa) | 79-80 | 0.322 ± 0.129 |
| 18 | m-s | Asn1(H)-Asn1(Hba) | 86-88 | 0.365 ± 0.146 |
| 19 | m-s | Ile6(H)-Ile6(Hg2*) | 74-112 | 0.306 ± 0.122 |
| 20 | m-s | Gly7(H)-Ile6(Hg2*) | 79-112 | 0.403 ± 0.161 |

##### (b) (continued) NOE distances for $2_M$.

| ID | type | name | atoms | $R^{\mathrm{NMR}}$ [nm] |
|---|---|---|---|---|
| 21 | m-s | Asn1(H)-Gly7(Hab) | 86-81 | 0.418 ± 0.167 |
| 22 | m-s | Asn1(H)-Cys8(Ha) | 86-83 | 0.365 ± 0.146 |
| 23 | m-s | Asn1(H)-Cys8(Hba) | 86-84 | 0.421 ± 0.168 |
| 24 | m-s | Asn1(H)-Hyp2(Hg) | 86-93 | 0.423 ± 0.169 |
| 25 | m-s | Ile6(H)-Trp4(Ha) | 74-65 | 0.41 ± 0.164 |
| 26 | s-s | Trp4(Hz2)-Trp4(He1) | 70-68 | 0.342 ± 0.137 |
| 27 | s-s | Cys8(Hbb)-Trp4(He1) | 85-68 | 0.307 ± 0.123 |
| 28 | s-s | Cys8(Hba)-Trp4(He1) | 84-68 | 0.383 ± 0.153 |
| 29 | s-s | Gly7(Hab)-Gly7(Haa) | 81-80 | 0.238 ± 0.095 |
| 30 | s-s | Gly7(Hab)-Gly7(Haa) | 81-80 | 0.256 ± 0.102 |
| 31 | s-s | Asn1(Ha)-Asn1(Hba) | 87-88 | 0.305 ± 0.122 |
| 32 | s-s | Asn1(Ha)-Asn1(Hba) | 87-88 | 0.319 ± 0.128 |
| 33 | s-s | Asn1(Ha)-Asn1(Hbb) | 87-89 | 0.347 ± 0.139 |
| 34 | s-s | Asn1(Hd2a)-Asn1(Hbb) | 115-89 | 0.367 ± 0.147 |
| 35 | s-s | Asn1(Hd2b)-Asn1(Hbb) | 116-89 | 0.338 ± 0.135 |
| 36 | s-s | Asn1(Hd2b)-Asn1(Hba) | 116-88 | 0.353 ± 0.141 |
| 37 | s-s | Asn1(Hd2a)-Asn1(Hba) | 115-88 | 0.393 ± 0.157 |
| 38 | s-s | Ile6(Hb)-Ile6(Ha) | 76-75 | 0.324 ± 0.13 |
| 39 | s-s | Hyp2(Hba)-Asn1(Ha) | 95-87 | 0.398 ± 0.159 |
| 40 | s-s | Hyp2(Hg)-Asn1(Ha) | 93-87 | 0.266 ± 0.107 |
| 41 | s-s | Ile3(Ha)-Asn1(Ha) | 97-87 | 0.337 ± 0.135 |

#### 2.3 Data from MD simulations of $M_{\text{ansa}}$- and $P_{\text{ansa}}$-Gly5Sar-amanullin

##### 2.3.1 $M_{\text{ansa}}$-Gly5Sar-amanullin

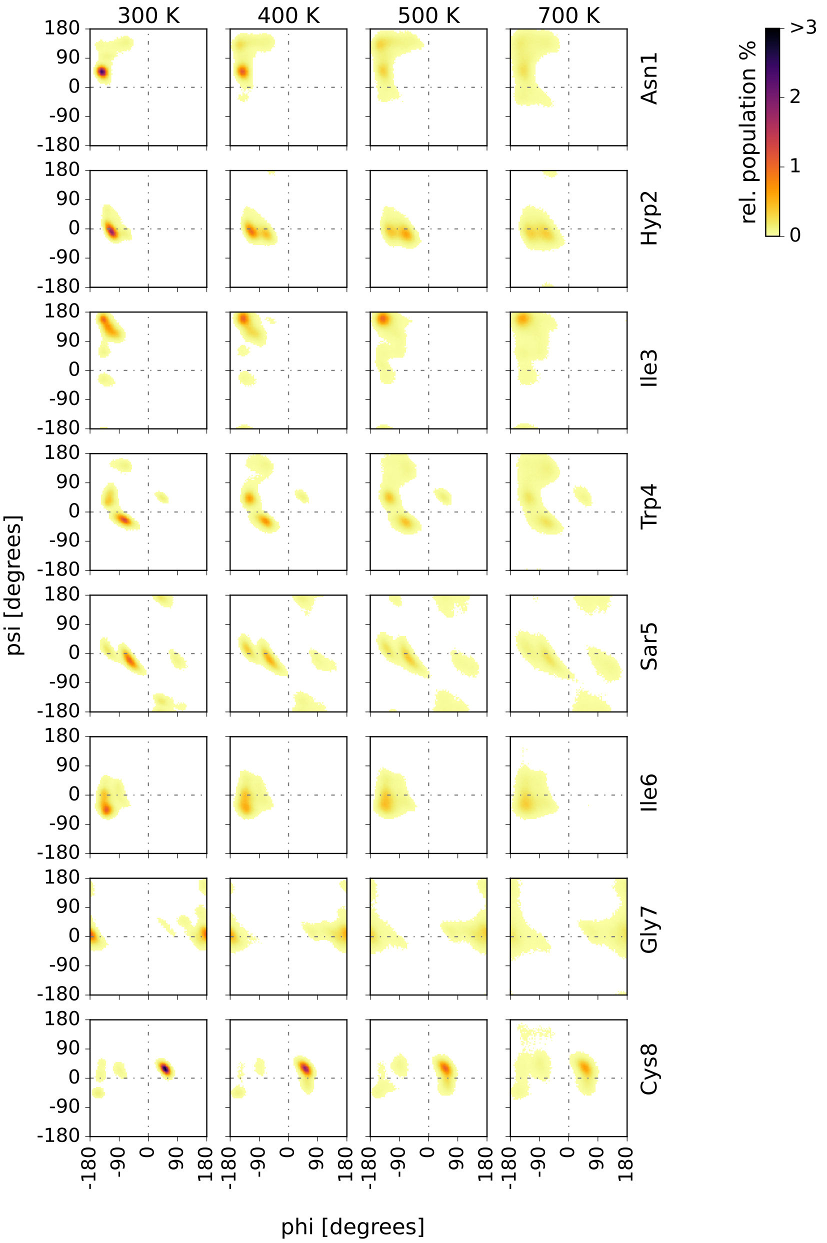

**Supplementary Figure 6.** Two-dimensional probability distributions of the $\phi$ and $\psi$ backbone torsion angles (Ramachandran plots) for the MD ensembles of $M_{\text{ansa}}$-Gly5Sar-amanullin at different temperatures. In each representation, the 2D-$(\phi, \psi)$-distribution is the average distribution over all 24 trajectories (in total 24 $\mu$s). For an overview of the simulated data, please refer to Supplementary Table 10.

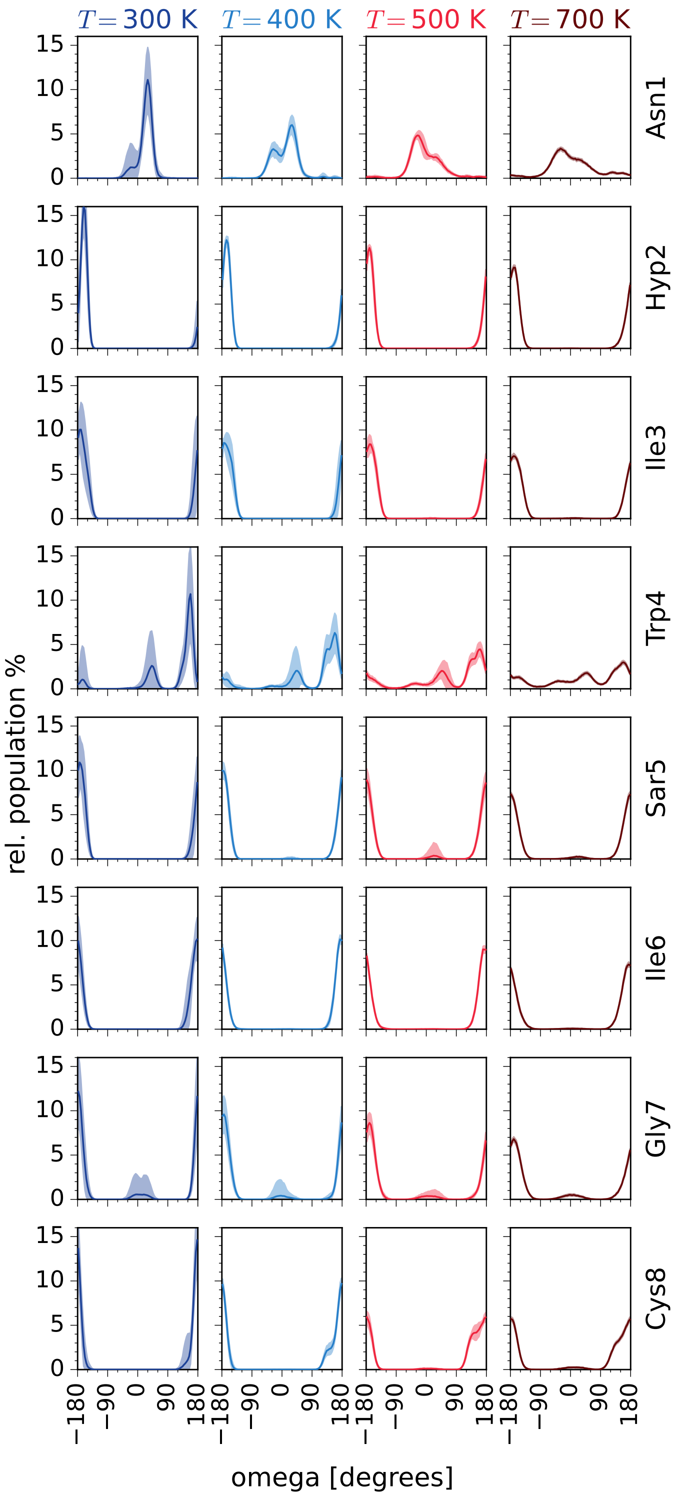

**Supplementary Figure 7.** Probability distributions of the residue-specific $\omega$-angles in the MD ensemble of $M_{\text{ansa}}$-Gly5Sar-amanullin at different temperatures. The distributions are shown as averages (solid line) and standard deviation (shaded area) over all 24 trajectories (in total 24 $\mu$s), respectively. For the definition of $\omega$, please refer to 3.2.2.

**Supplementary Table 5.** Configurations of the $\omega$-torsion angles of $M_{\text{ansa}}$-Gly5Sar-amanullin at 300 K broken down by individual trajectories ('Traj'). Populations of minor configurations are shown in brackets. Based on the configurations of the $\omega$-torsion angles of Trp4 and Gly7, all trajectories were assigned to one of the following four sub data sets ('Set'): 'tt' : trans, trans; 'tc' : trans, cis; 'ct' : cis, trans; 'cc' : cis, cis, where the first configuration describes $\omega$(Trp4) and the second $\omega$(Gly7). For the definition of $\omega$, please refer to 3.2.2.

| Traj | Set | $\omega$(Asn1) | $\omega$(Hyp2) | $\omega$(Ile3) | $\omega$(Trp4) | $\omega$(Sar5) | $\omega$(Ile6) | $\omega$(Gly7) | $\omega$(Cys8) |
|---|---|---|---|---|---|---|---|---|---|
| 1 | tt | cis | trans | trans | trans | trans | trans | trans | trans |
| 2 | tt | cis (trans $<$ 1%) | trans | trans | trans (cis: 15%) | trans | trans | trans | trans |
| 3 | tt | cis (trans $<$ 1%) | trans | trans | trans (cis: 33%) | trans | trans | trans | trans |
| 4 | tt | cis (trans $<$ 1%) | trans | trans | trans (cis: 8%) | trans | trans | trans | trans |
| 5 | tt | cis | trans | trans | trans (cis: 10%) | trans | trans | trans | trans |
| 6 | tt | cis | trans | trans | trans (cis: 15%) | trans | trans | trans | trans |
| 7 | tt | cis | trans | trans | trans (cis: 11%) | trans | trans | trans | trans |
| 8 | ct | cis | trans | trans | cis (trans: 1%) | trans | trans | trans | trans |
| 9 | tt | cis (trans $<$ 1%) | trans | trans | trans (cis: 29%) | trans | trans | trans | trans |
| 10 | tt | cis | trans | trans | trans (cis: 26%) | trans | trans | trans | trans |
| 11 | tt | cis | trans | trans | trans (cis: 1%) | trans | trans | trans | trans |
| 12 | tt | cis | trans | trans | trans (cis: 3%) | trans | trans | trans | trans |
| 13 | tt | cis (trans $<$ 1%) | trans | trans | trans (cis: 1%) | trans | trans | trans | trans |
| 14 | tt | cis (trans $<$ 1%) | trans | trans | trans | trans | trans | trans | trans |
| 15 | tt | cis | trans | trans | trans (cis: 5%) | trans | trans | trans | trans |
| 16 | tt | cis (trans $<$ 1%) | trans | trans | trans (cis: 6%) | trans | trans | trans | trans |
| 17 | tt | cis (trans $<$ 1%) | trans | trans | trans (cis: 3%) | trans | trans | trans | trans |
| 18 | tt | cis | trans | trans | trans (cis: 11%) | trans | trans | trans | trans |
| 19 | tt | cis | trans | trans | trans (cis: 10%) | trans | trans | trans | trans |
| 20 | ct | cis | trans | trans | cis | trans | trans | trans | trans |
| 21 | tt | cis | trans | trans | trans (cis: 11%) | trans | trans | trans | trans |
| 22 | tc | cis | trans | trans | trans | trans | trans | cis | trans |
| 23 | tt | cis (trans $<$ 1%) | trans | trans | trans (cis: 18%) | trans | trans | trans | trans |
| 24 | cc | cis | trans | trans | cis | trans | trans | cis | trans |

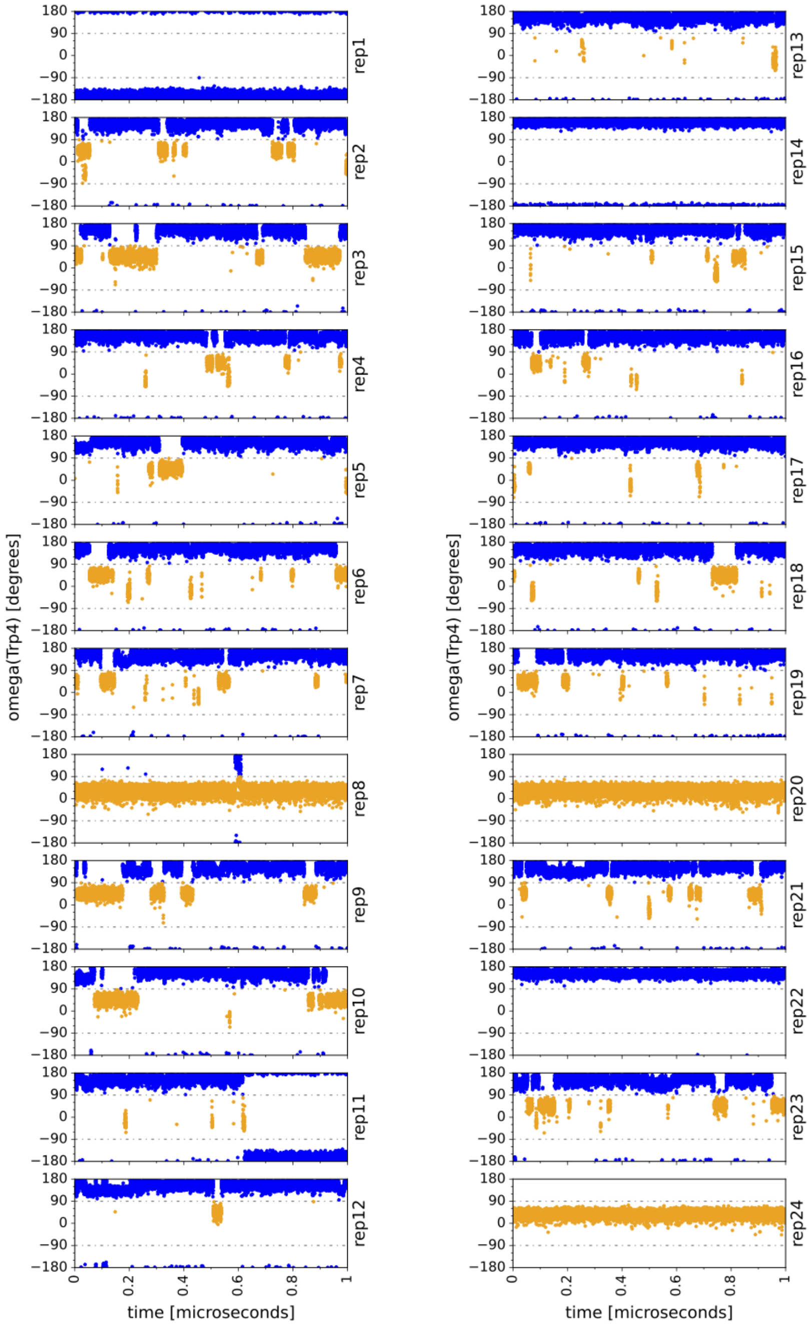

**Supplementary Figure 8.** Time series for the $\omega$(Trp4) torsion angle shown for all 24 replicas of $M_{\mathrm{ansa}}$-Gly5Sar-amanullin at 300 K (in total 24 $\mu$s). To distinguish the 'cis' (orange) and 'trans' (blue) states, the threshold $\pm 90\,^\circ$ (grey line) was used. For the definition of $\omega$, please refer to 3.2.2.

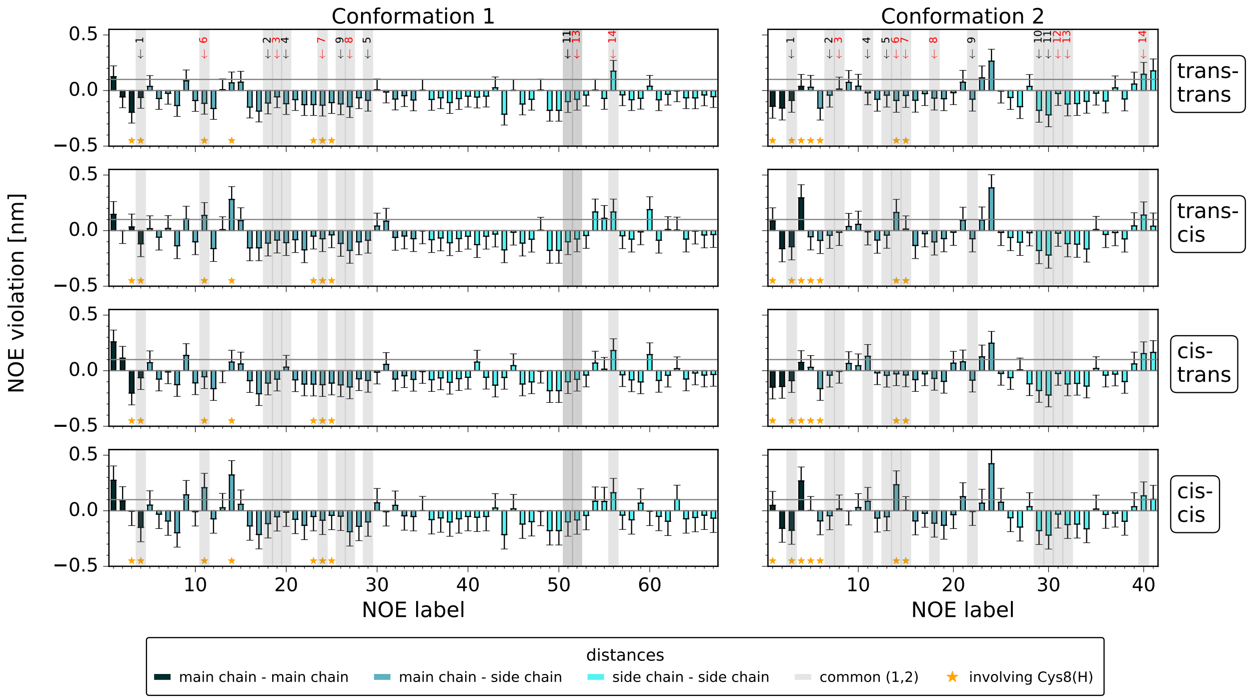

**Supplementary Figure 9.** Differences in the NOE upper distance bounds and the distances in the MD ensemble, calculated for the NOE sets of $1_M$ and $2_M$ and the four sub data sets of the MD ensemble of $M_{\text{ansa}}$-Gly5Sar-amanullin at 300 K. Each sub data set refers to one of the following four ($\omega$(Trp4), $\omega$(Gly7))-configurations as indicated on the right: (trans, trans), (trans,cis), (cis, trans) and (cis,cis). For the assignment of the trajectories to these sub data sets, please refer to Supplementary Table 5.

The distances from MD were calculated as averages over all frames of each sub data set, respectively (see 3.2.4, eq. 4). Three different types of distances are distinguished according to the assignment of the protons to the main chain ('m'; atoms: $H_N$), the side chain ('s'; atoms: else) or (only for Sar5) the main chain II ('x'; H of the methyl group replacing the $H_N$): 'm-m' or 'm-x' : blue; 'm-s', 'x-s' : light blue, 's-s' : cyan. If the difference between NMR and MD is larger than 0.1 nm (grey line), the distance is considered 'violated'. Inter-proton distances that could be obtained for both conformations are highlighted in grey and numbered from 1 to 14. Red numbers indicate that the difference between $1_M$ and $2_M$ is larger than 0.05 nm. Distances involving the amide $H_N$ of Cys8 are indicated by orange stars. For an overview of the inter proton distances from NMR, please refer to Supplementary Table 4. For the associated values from MD, please refer to Supplementary Table 6.

**Supplementary Table 6.** Inter-proton distances calculated as averages ± standard deviations relative to the size of the sub data sets of the MD ensemble of $M_{\text{ansa}}$-Gly5Sar-amanullin at 300 K. The distances were calculated for the atom pairs defined in Supplementary Table 4. Each sub data set refers to one of the following four ($\omega$(Trp4), $\omega$(Gly7))-configurations as indicated by the indices: 'tt' : (trans, trans), 'tc' : (trans,cis), 'ct' : (cis, trans) and 'cc' : (cis,cis). For the assignment of the trajectories to these sub data sets, please refer to Supplementary Table 5. For the calculation details, please refer to 3.2.4.

**(a)** NOE distances referring to $1_M$.

| ID | $R^{\text{tt}}$ [nm] | $R^{\text{tc}}$ [nm] | $R^{\text{ct}}$ [nm] | $R^{\text{cc}}$ [nm] |
|---|---|---|---|---|
| 1 | 0.125 ± 0.098 | 0.146 ± 0.116 | 0.259 ± 0.106 | 0.274 ± 0.129 |
| 2 | -0.057 ± 0.098 | -0.001 ± 0.116 | 0.113 ± 0.106 | 0.088 ± 0.129 |
| 3 | -0.195 ± 0.098 | 0.033 ± 0.116 | -0.201 ± 0.106 | -0.005 ± 0.129 |
| 4 | -0.062 ± 0.098 | -0.119 ± 0.116 | -0.063 ± 0.106 | -0.15 ± 0.129 |
| 5 | 0.037 ± 0.098 | 0.018 ± 0.116 | 0.072 ± 0.106 | 0.051 ± 0.129 |
| 6 | -0.071 ± 0.098 | -0.059 ± 0.116 | -0.076 ± 0.106 | -0.031 ± 0.129 |
| 7 | -0.025 ± 0.098 | 0.02 ± 0.116 | -0.011 ± 0.106 | -0.092 ± 0.129 |
| 8 | -0.135 ± 0.098 | -0.136 ± 0.116 | -0.126 ± 0.106 | -0.198 ± 0.129 |
| 9 | 0.088 ± 0.098 | 0.104 ± 0.116 | 0.137 ± 0.106 | 0.143 ± 0.129 |
| 10 | -0.09 ± 0.098 | -0.101 ± 0.116 | -0.11 ± 0.106 | -0.008 ± 0.129 |
| 11 | -0.115 ± 0.098 | 0.137 ± 0.116 | -0.055 ± 0.106 | 0.208 ± 0.129 |
| 12 | -0.162 ± 0.098 | -0.162 ± 0.116 | -0.162 ± 0.106 | -0.073 ± 0.129 |
| 13 | 0.008 ± 0.098 | 0.009 ± 0.116 | -0.002 ± 0.106 | 0.027 ± 0.129 |
| 14 | 0.069 ± 0.098 | 0.281 ± 0.116 | 0.078 ± 0.106 | 0.322 ± 0.129 |
| 15 | 0.076 ± 0.098 | 0.09 ± 0.116 | 0.061 ± 0.106 | 0.058 ± 0.129 |
| 16 | -0.149 ± 0.098 | -0.155 ± 0.116 | -0.09 ± 0.106 | -0.136 ± 0.129 |
| 17 | -0.185 ± 0.098 | -0.155 ± 0.116 | -0.208 ± 0.106 | -0.214 ± 0.129 |
| 18 | -0.112 ± 0.098 | -0.112 ± 0.116 | -0.112 ± 0.106 | -0.116 ± 0.129 |
| 19 | -0.056 ± 0.098 | -0.085 ± 0.116 | -0.076 ± 0.106 | -0.053 ± 0.129 |
| 20 | -0.118 ± 0.098 | -0.108 ± 0.116 | 0.032 ± 0.106 | -0.013 ± 0.129 |
| 21 | -0.082 ± 0.098 | -0.082 ± 0.116 | -0.082 ± 0.106 | -0.077 ± 0.129 |
| 22 | -0.127 ± 0.098 | -0.173 ± 0.116 | -0.12 ± 0.106 | -0.131 ± 0.129 |
| 23 | -0.126 ± 0.098 | -0.05 ± 0.116 | -0.121 ± 0.106 | -0.052 ± 0.129 |
| 24 | -0.132 ± 0.098 | -0.073 ± 0.116 | -0.126 ± 0.106 | -0.085 ± 0.129 |
| 25 | -0.108 ± 0.098 | -0.038 ± 0.116 | -0.111 ± 0.106 | -0.041 ± 0.129 |
| 26 | -0.12 ± 0.098 | -0.115 ± 0.116 | -0.128 ± 0.106 | -0.049 ± 0.129 |
| 27 | -0.145 ± 0.098 | -0.178 ± 0.116 | -0.146 ± 0.106 | -0.188 ± 0.129 |
| 28 | -0.065 ± 0.098 | -0.101 ± 0.116 | -0.073 ± 0.106 | -0.14 ± 0.129 |
| 29 | -0.088 ± 0.098 | -0.087 ± 0.116 | -0.088 ± 0.106 | -0.1 ± 0.129 |
| 30 | 0.007 ± 0.098 | 0.041 ± 0.116 | -0.013 ± 0.106 | 0.071 ± 0.129 |
| 31 | -0.013 ± 0.098 | 0.085 ± 0.116 | 0.057 ± 0.106 | -0.008 ± 0.129 |
| 32 | -0.077 ± 0.098 | -0.061 ± 0.116 | -0.075 ± 0.106 | 0.049 ± 0.129 |
| 33 | -0.046 ± 0.098 | -0.048 ± 0.116 | -0.046 ± 0.106 | -0.045 ± 0.129 |

**(a)** (continued) NOE distances referring to $1_M$.

| ID | $R^{\text{tt}}$ [nm] | $R^{\text{tc}}$ [nm] | $R^{\text{ct}}$ [nm] | $R^{\text{cc}}$ [nm] |
|---|---|---|---|---|
| 34 | -0.085 ± 0.098 | -0.073 ± 0.116 | -0.078 ± 0.106 | -0.052 ± 0.129 |
| 35 | 0.003 ± 0.098 | -0.003 ± 0.116 | -0.008 ± 0.106 | 0.0 ± 0.129 |
| 36 | -0.08 ± 0.098 | -0.082 ± 0.116 | -0.081 ± 0.106 | -0.08 ± 0.129 |
| 37 | -0.065 ± 0.098 | -0.066 ± 0.116 | -0.065 ± 0.106 | -0.064 ± 0.129 |
| 38 | -0.108 ± 0.098 | -0.109 ± 0.116 | -0.105 ± 0.106 | -0.101 ± 0.129 |
| 39 | -0.072 ± 0.098 | -0.072 ± 0.116 | -0.072 ± 0.106 | -0.072 ± 0.129 |
| 40 | -0.051 ± 0.098 | -0.051 ± 0.116 | -0.06 ± 0.106 | -0.052 ± 0.129 |
| 41 | -0.059 ± 0.098 | -0.126 ± 0.116 | 0.077 ± 0.106 | -0.058 ± 0.129 |
| 42 | -0.051 ± 0.098 | -0.051 ± 0.116 | -0.06 ± 0.106 | -0.052 ± 0.129 |
| 43 | 0.025 ± 0.098 | -0.031 ± 0.116 | -0.13 ± 0.106 | 0.025 ± 0.129 |
| 44 | -0.213 ± 0.098 | -0.173 ± 0.116 | -0.061 ± 0.106 | -0.215 ± 0.129 |
| 45 | 0.002 ± 0.098 | -0.015 ± 0.116 | 0.045 ± 0.106 | 0.018 ± 0.129 |
| 46 | -0.121 ± 0.098 | -0.112 ± 0.116 | -0.12 ± 0.106 | -0.12 ± 0.129 |
| 47 | -0.079 ± 0.098 | -0.078 ± 0.116 | -0.101 ± 0.106 | -0.098 ± 0.129 |
| 48 | 0.005 ± 0.098 | 0.004 ± 0.116 | -0.005 ± 0.106 | -0.006 ± 0.129 |
| 49 | -0.178 ± 0.098 | -0.178 ± 0.116 | -0.18 ± 0.106 | -0.179 ± 0.129 |
| 50 | -0.178 ± 0.098 | -0.178 ± 0.116 | -0.18 ± 0.106 | -0.179 ± 0.129 |
| 51 | -0.1 ± 0.098 | -0.1 ± 0.116 | -0.1 ± 0.106 | -0.1 ± 0.129 |
| 52 | -0.079 ± 0.098 | -0.075 ± 0.116 | -0.078 ± 0.106 | -0.082 ± 0.129 |
| 53 | -0.041 ± 0.098 | -0.044 ± 0.116 | -0.039 ± 0.106 | -0.042 ± 0.129 |
| 54 | 0.004 ± 0.098 | 0.168 ± 0.116 | 0.069 ± 0.106 | 0.085 ± 0.129 |
| 55 | -0.071 ± 0.098 | 0.11 ± 0.116 | 0.014 ± 0.106 | 0.083 ± 0.129 |
| 56 | 0.174 ± 0.098 | 0.168 ± 0.116 | 0.181 ± 0.106 | 0.162 ± 0.129 |
| 57 | -0.039 ± 0.098 | -0.046 ± 0.116 | -0.037 ± 0.106 | -0.039 ± 0.129 |
| 58 | -0.082 ± 0.098 | -0.077 ± 0.116 | -0.083 ± 0.106 | -0.081 ± 0.129 |
| 59 | -0.07 ± 0.098 | -0.011 ± 0.116 | -0.101 ± 0.106 | 0.069 ± 0.129 |
| 60 | 0.039 ± 0.098 | 0.189 ± 0.116 | 0.145 ± 0.106 | -0.005 ± 0.129 |
| 61 | -0.083 ± 0.098 | -0.087 ± 0.116 | -0.088 ± 0.106 | -0.051 ± 0.129 |
| 62 | -0.033 ± 0.098 | 0.01 ± 0.116 | -0.041 ± 0.106 | -0.094 ± 0.129 |
| 63 | -0.003 ± 0.098 | 0.006 ± 0.116 | 0.018 ± 0.106 | 0.101 ± 0.129 |
| 64 | -0.065 ± 0.098 | -0.076 ± 0.116 | -0.075 ± 0.106 | -0.072 ± 0.129 |
| 65 | -0.066 ± 0.098 | -0.004 ± 0.116 | -0.021 ± 0.106 | -0.06 ± 0.129 |
| 66 | -0.04 ± 0.098 | -0.039 ± 0.116 | -0.039 ± 0.106 | -0.04 ± 0.129 |
| 67 | -0.055 ± 0.098 | -0.036 ± 0.116 | -0.034 ± 0.106 | -0.067 ± 0.129 |

**Supplementary Table 6.** (continued).

**(b)** NOE distances referring to $2_M$.

| ID | $R^{tt}$ [nm] | $R^{tc}$ [nm] | $R^{ct}$ [nm] | $R^{cc}$ [nm] |
|---|---|---|---|---|
| 1 | -0.142 ± 0.107 | 0.087 ± 0.118 | -0.148 ± 0.106 | 0.049 ± 0.125 |
| 2 | -0.158 ± 0.107 | -0.162 ± 0.118 | -0.142 ± 0.106 | -0.158 ± 0.125 |
| 3 | -0.088 ± 0.107 | -0.145 ± 0.118 | -0.09 ± 0.106 | -0.176 ± 0.125 |
| 4 | 0.038 ± 0.107 | 0.295 ± 0.118 | 0.074 ± 0.106 | 0.269 ± 0.125 |
| 5 | 0.029 ± 0.107 | -0.055 ± 0.118 | 0.03 ± 0.106 | 0.003 ± 0.125 |
| 6 | -0.158 ± 0.107 | -0.088 ± 0.118 | -0.161 ± 0.106 | -0.091 ± 0.125 |
| 7 | -0.042 ± 0.107 | -0.042 ± 0.118 | -0.041 ± 0.106 | -0.045 ± 0.125 |
| 8 | 0.014 ± 0.107 | -0.015 ± 0.118 | -0.006 ± 0.106 | 0.017 ± 0.125 |
| 9 | 0.073 ± 0.107 | 0.037 ± 0.118 | 0.065 ± 0.106 | -0.002 ± 0.125 |
| 10 | 0.039 ± 0.107 | 0.056 ± 0.118 | 0.045 ± 0.106 | 0.029 ± 0.125 |
| 11 | -0.021 ± 0.107 | -0.011 ± 0.118 | 0.129 ± 0.106 | 0.085 ± 0.125 |
| 12 | -0.079 ± 0.107 | -0.085 ± 0.118 | -0.02 ± 0.106 | -0.066 ± 0.125 |
| 13 | -0.042 ± 0.107 | -0.04 ± 0.118 | -0.042 ± 0.106 | -0.054 ± 0.125 |
| 14 | -0.089 ± 0.107 | 0.163 ± 0.118 | -0.028 ± 0.106 | 0.234 ± 0.125 |
| 15 | -0.044 ± 0.107 | 0.014 ± 0.118 | -0.039 ± 0.106 | 0.002 ± 0.125 |
| 16 | -0.087 ± 0.107 | -0.133 ± 0.118 | -0.081 ± 0.106 | -0.092 ± 0.125 |
| 17 | -0.028 ± 0.107 | -0.028 ± 0.118 | -0.028 ± 0.106 | -0.023 ± 0.125 |
| 18 | -0.068 ± 0.107 | -0.101 ± 0.118 | -0.07 ± 0.106 | -0.112 ± 0.125 |
| 19 | -0.074 ± 0.107 | -0.072 ± 0.118 | -0.096 ± 0.106 | -0.131 ± 0.125 |
| 20 | -0.019 ± 0.107 | -0.015 ± 0.118 | 0.068 ± 0.106 | -0.035 ± 0.125 |

**(b)** NOE distances referring to $2_M$.

| ID | $R^{tt}$ [nm] | $R^{tc}$ [nm] | $R^{ct}$ [nm] | $R^{cc}$ [nm] |
|---|---|---|---|---|
| 21 | 0.075 ± 0.107 | 0.092 ± 0.118 | 0.08 ± 0.106 | 0.127 ± 0.125 |
| 22 | -0.078 ± 0.107 | -0.073 ± 0.118 | -0.085 ± 0.106 | -0.007 ± 0.125 |
| 23 | 0.114 ± 0.107 | 0.094 ± 0.118 | 0.125 ± 0.106 | 0.068 ± 0.125 |
| 24 | 0.265 ± 0.107 | 0.384 ± 0.118 | 0.248 ± 0.106 | 0.423 ± 0.125 |
| 25 | -0.005 ± 0.107 | -0.016 ± 0.118 | -0.025 ± 0.106 | 0.076 ± 0.125 |
| 26 | -0.063 ± 0.107 | -0.062 ± 0.118 | -0.062 ± 0.106 | -0.063 ± 0.125 |
| 27 | -0.145 ± 0.107 | -0.105 ± 0.118 | 0.007 ± 0.106 | -0.147 ± 0.125 |
| 28 | 0.037 ± 0.107 | -0.019 ± 0.118 | -0.118 ± 0.106 | 0.037 ± 0.125 |
| 29 | -0.178 ± 0.107 | -0.179 ± 0.118 | -0.178 ± 0.106 | -0.179 ± 0.125 |
| 30 | -0.219 ± 0.107 | -0.219 ± 0.118 | -0.219 ± 0.106 | -0.219 ± 0.125 |
| 31 | -0.027 ± 0.107 | -0.023 ± 0.118 | -0.025 ± 0.106 | -0.03 ± 0.125 |
| 32 | -0.121 ± 0.107 | -0.117 ± 0.118 | -0.119 ± 0.106 | -0.124 ± 0.125 |
| 33 | -0.117 ± 0.107 | -0.122 ± 0.118 | -0.114 ± 0.106 | -0.12 ± 0.125 |
| 34 | -0.097 ± 0.107 | -0.165 ± 0.118 | -0.138 ± 0.106 | -0.162 ± 0.125 |
| 35 | -0.024 ± 0.107 | 0.01 ± 0.118 | 0.02 ± 0.106 | 0.015 ± 0.125 |
| 36 | -0.094 ± 0.107 | -0.034 ± 0.118 | -0.04 ± 0.106 | -0.033 ± 0.125 |
| 37 | 0.024 ± 0.107 | -0.018 ± 0.118 | -0.031 ± 0.106 | -0.021 ± 0.125 |
| 38 | -0.076 ± 0.107 | -0.074 ± 0.118 | -0.098 ± 0.106 | -0.095 ± 0.125 |
| 39 | 0.059 ± 0.107 | 0.043 ± 0.118 | 0.061 ± 0.106 | 0.037 ± 0.125 |
| 40 | 0.146 ± 0.107 | 0.14 ± 0.118 | 0.153 ± 0.106 | 0.134 ± 0.125 |
| 41 | 0.177 ± 0.107 | 0.04 ± 0.118 | 0.164 ± 0.106 | 0.103 ± 0.125 |

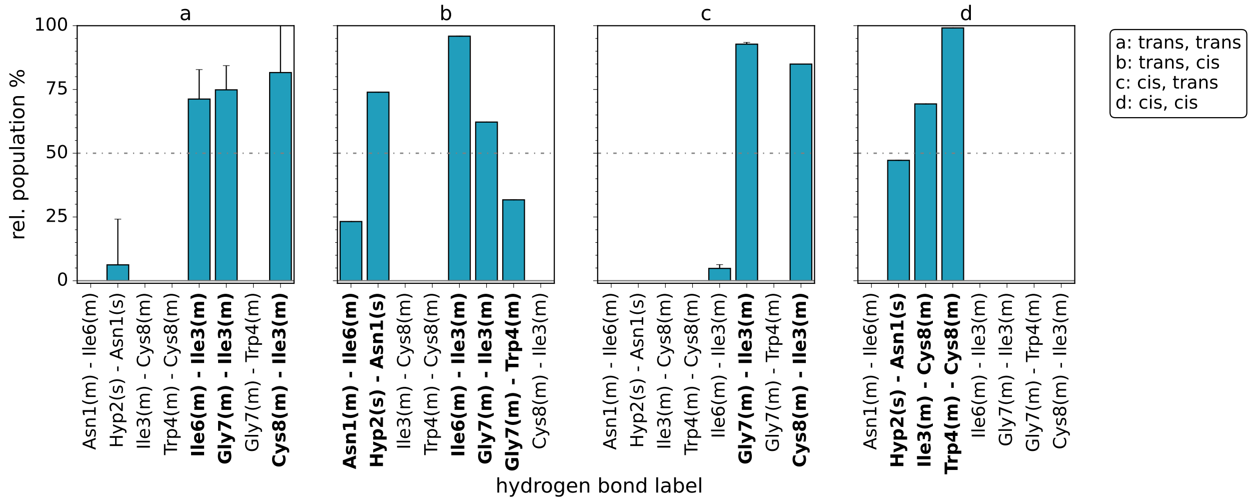

**Supplementary Figure 10.** Hydrogen bond populations (bars) for the four different sub data sets of $M_{\mathrm{ansa}}$-Gly5Sar-amanullin at 300 K. Each sub data set refers to one of the following four ($\omega$(Trp4), $\omega$(Gly7))-configurations: a : (trans, trans); b : (trans, cis); c : (cis, trans); d : (cis, cis).
Hydrogen bonds are indicated that occur in at least 20% of the structures of at least one sub data set (a-d). For each sub data set, the hydrogen bonds with a relative occurrence ≥20% are additionally highlighted with bold font. All hydrogen bonds are labelled as 'donor - acceptor', and differentiated in main chain ('m', atoms: N, $H_N$, $C_\alpha$, $C_{CO}$ and $O_{CO}$) and side chain ('s', atoms: else) interactions. Please refer to 3.2.1 for further details on the computation of the hydrogen bonds.

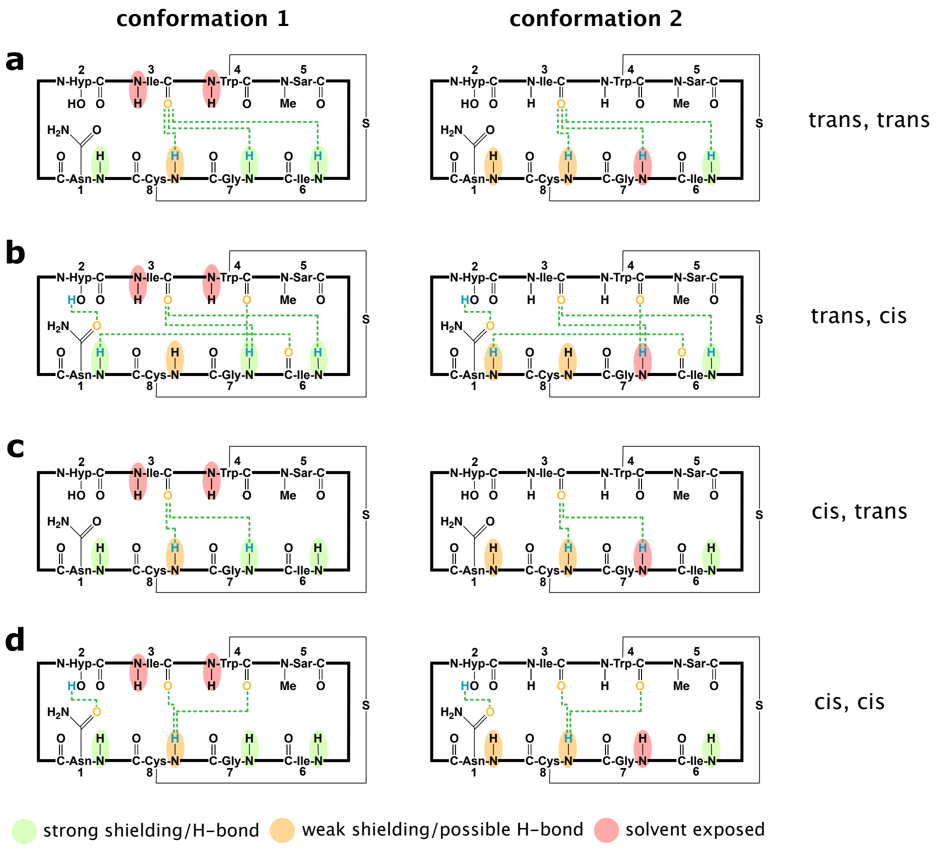

**Supplementary Figure 11.** Hydrogen bond schemes showing the hydrogen bonds with an occurrence ≥20% in the four sub data sets of the MD ensemble of $M_{\text{ansa}}$-Gly5Sar-amanullin at 300 K. Each sub data set comprises one of the following four ($\omega$(Trp4), $\omega$(Gly7))-configurations: a: (trans, trans); b: (trans, cis); c: (cis, trans); d: (cis, cis). The hydrogen bond populations were always calculated in relation to the size of the respective sub data set. Based on the temperature coefficients obtained for $1_M$ and $2_M$ (Supplementary Table 2), the amides are coloured according to their shielding from solvent: left : $1_M$; right : $2_M$; red : solvent exposed; orange : weak shielding, possibly involved in hydrogen bonding; green : strong shielding or hydrogen bonding.

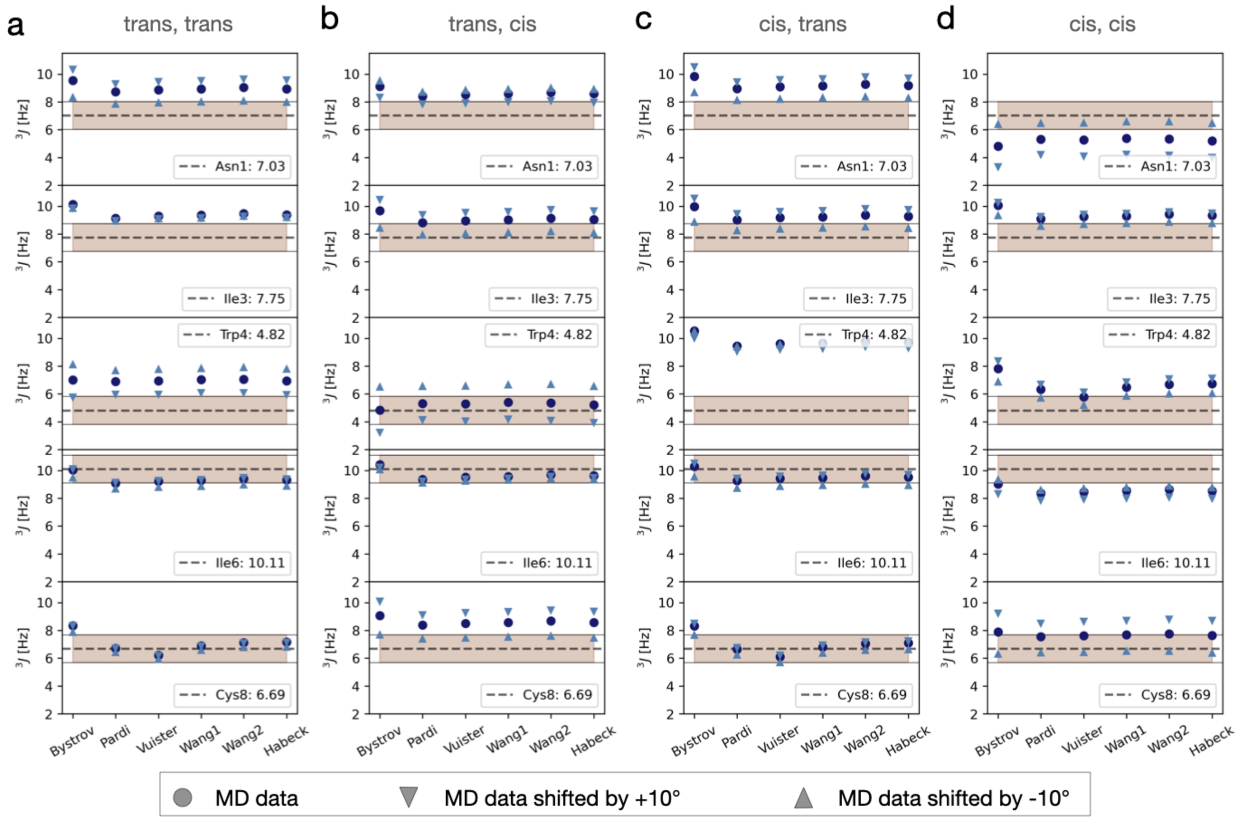

**Supplementary Figure 12.** $^3J_{\mathrm{H^N-H^\alpha}}$-coupling constants for the four different sub data sets (a-d) of $M_{\mathrm{ansa}}$-Gly5Sar-amanullin at 300 K. Each sub data set denotes one of the following four ($\omega$(Trp4), $\omega$(Gly7)-configurations: **a:** (trans, trans); **b:** (trans, cis); **c:** (cis, trans); **d:** (cis, cis). For the assignment of the trajectories to these sub data sets, please refer to Supplementary Table 5.
*dashed line and brown area:* experimental value (Supplementary Table 3) and ±1 Hz uncertainty. To account for force-field inaccuracies, we additionally shifted the $\phi_{\mathrm{H^N H^\alpha}}$-trajectories by +10° and -10° and recalculated the $^3J_{\mathrm{H^N-H^\alpha}}$-coupling constants (triangles). We considered six parametrisations of the Karplus curve: Bystrov,[5] Pardi,[6] Vuister,[7] Wang1,[3] Wang2,[3] and Habeck.[8]

**Supplementary Table 7.** Overview of the parameters used in Supplementary Figure 12 for the Karplus' equation ${}^3J_{\mathrm{H^N-H^\alpha}} = A \cdot \cos^2(\phi + \theta) + B \cdot \cos(\phi + \theta) + C$ to obtain ${}^3J_{\mathrm{H^N-H^\alpha}}$-coupling constants from MD data.

| Ref. | label | $A$ [Hz] | $B$ [Hz] | $C$ [Hz] | $\theta$ [°] |
|---|---|---|---|---|---|
| 3 | W1 | 6.64 | -1.43 | 1.86 | -60 |
| 3 | W2 | 6.98 | -1.38 | 1.72 | -60 |
| 5 | B | 9.4 | -1.1 | 0.4 | -60 |
| 6 | P | 6.4 | -1.4 | 1.9 | -60 |
| 7 | V | 6.51 | -1.76 | 1.6 | -60 |
| 8 | H | 7.13 | 1.31 | 1.56 | -60 |

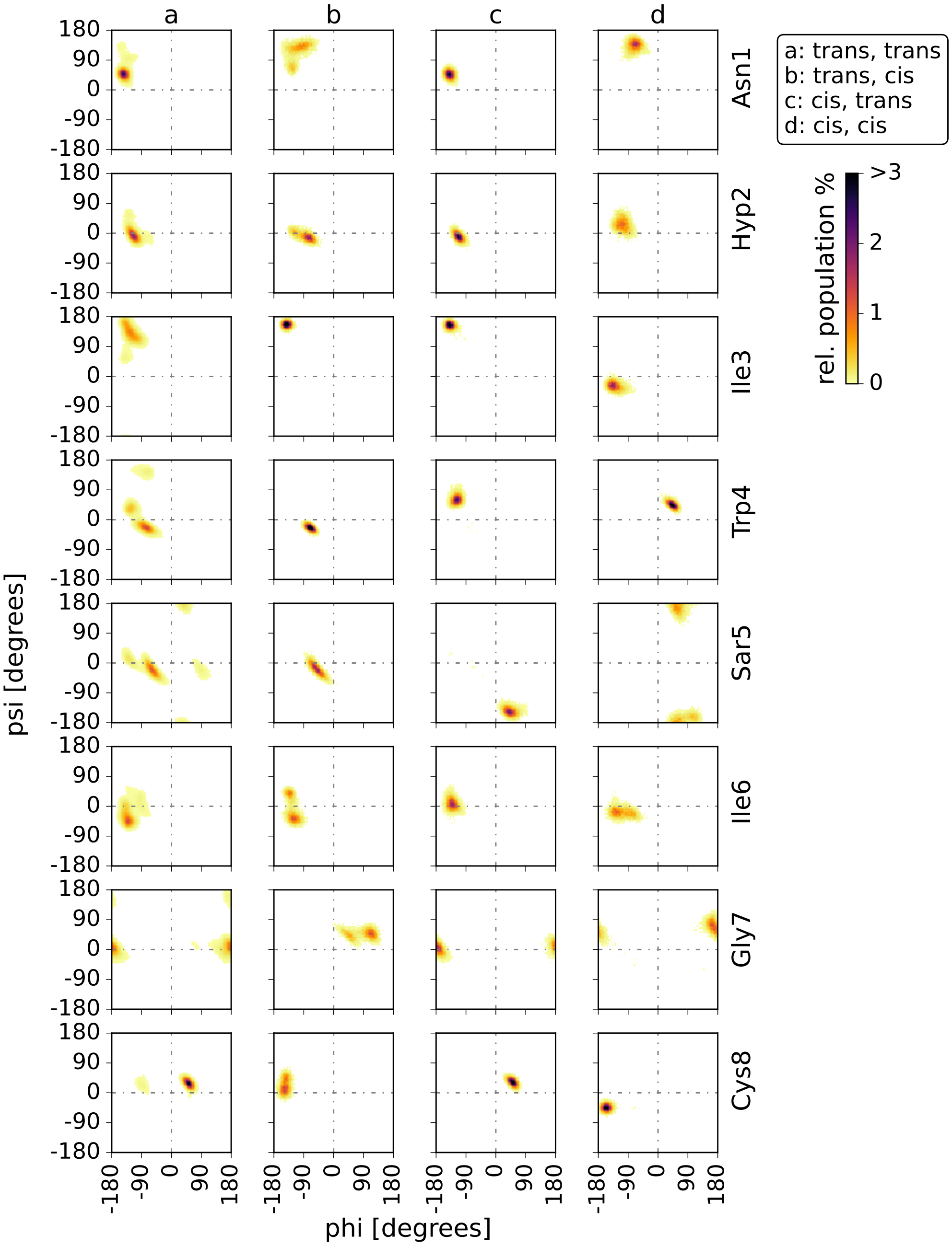

**Supplementary Figure 13.** Two-dimensional probability distributions of the $\phi$- and $\psi$-backbone torsion angles (Ramachandran plots) for the four different sub data sets of the MD ensemble of $M_{\text{ansa}}$-Gly5Sar-amanullin at 300 K. Each sub data set refers to one of the following four ($\omega$(Trp4), $\omega$(Gly7))-configurations: **a:** (trans, trans); **b:** (trans, cis); **c:** (cis, trans); **d:** (cis, cis).

The 2D-distributions shown here always represent the average distribution over all trajectories of each sub data set (a-d). For the assignment of the trajectories to these four sub data sets, please refer to Supplementary Table 5.

**Supplementary Table 8.** Radius cut-off $R$, neighbour cut-off $C$, minimal cluster size $M$ and noise $N\%$ for the clustering of the 'tt' sub data set of the $M_{\text{ansa}}$-Gly5Sar-amanullin MD ensemble at 300 K. Prior to the clustering, a discretisation was performed. For details on the discretisation method, please refer to 3.2.5. The input data set for the clustering comprised 200020 data points with four dimensions (time-independent components: 'TICs'). The clustering hierarchy level is denoted 'L', and the splitting of the clusters is provided ('split').

| L | $R$ | $C$ | $M$ | $N\%$ | split |
|---|---|---|---|---|---|
| 0 | 0.2 | 8 | 100 | 4.6 | 0 → 1,2,3,4,5 |

##### 2.3.2 $M_{\mathrm{ansa}}$-Gly5Sar-amanullin: *(trans-trans)*-configuration

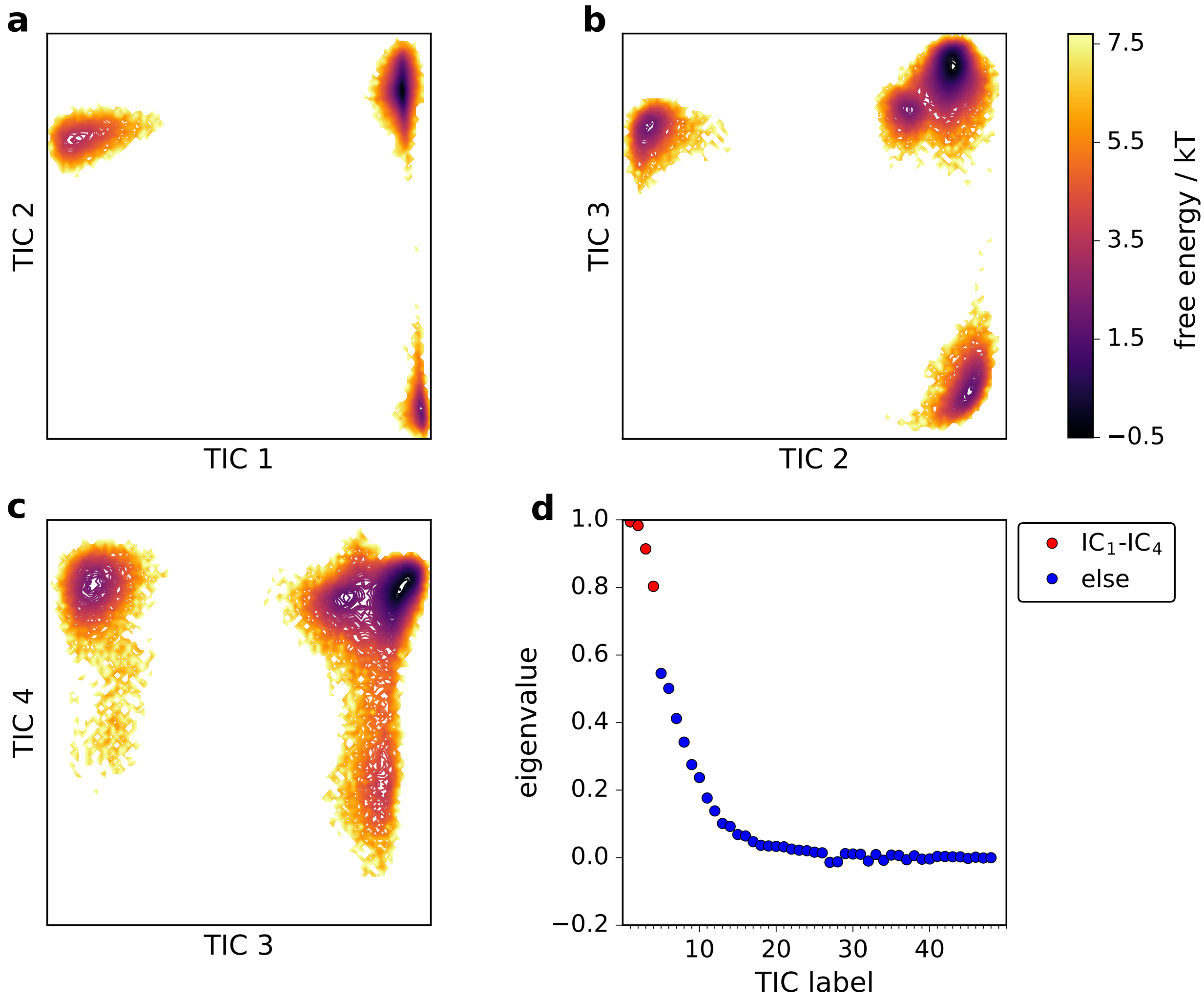

**Supplementary Figure 14.** a)-c) Projection of the 'tt' sub data set of 300 K MD ensemble of $M_{\mathrm{ansa}}$-Gly5Sar-amanullin on the first four time-independent components (TIC 1 - TIC 4). For each pair (a-c), a histogram was computed with a discretisation into 100 equally-sized bins for each component. The energy profiles for these [100 x 100]-histograms were then calculated by $F_{i,j} = -\ln(z_{i,j})$ where $z_{i,j}$ is the number of counts of cell $ij$. d) Eigenvalue spectrum for the eigenvector matrix containing the time-independent components. The eigenvalues are shown as dots, and for the first four time-independent components they are highlighted in red. For details on the TICA method and the parameters used, please refer to 3.2.5.

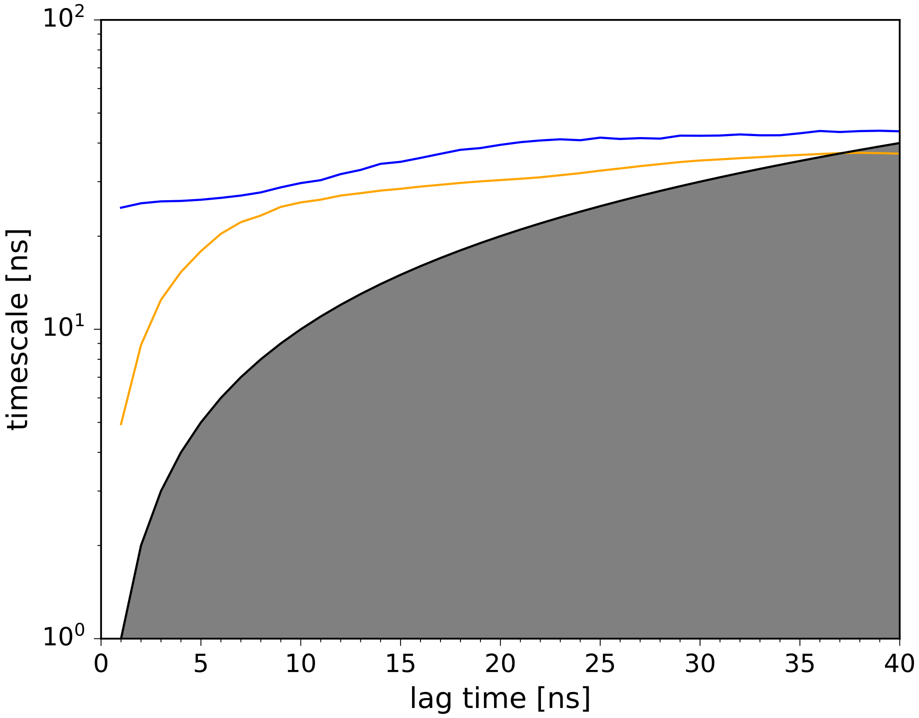

**Supplementary Figure 15.** Implied timescales ('timescales') from core-set Markov state models (cs-MSM) estimated at different lag times ('lag time'). The implied timescales are coloured according to the eigenvectors, which represent the slowest processes in the cs-MSM: blue : 2nd eigenvector (slowest process), orange : 3rd eigenvector (2nd slowest process). The value range, in which the timescales become unphysical is shaded. The black line represents 'lag time' = 'timescale'. The cs-MSMs were constructed on the clustered 'tt' sub data set of the $M_{\text{ansa}}$-Gly5Sar-amanullin MD ensemble at 300 K. For further details on the cs-MSM construction, please refer to 3.2.7.

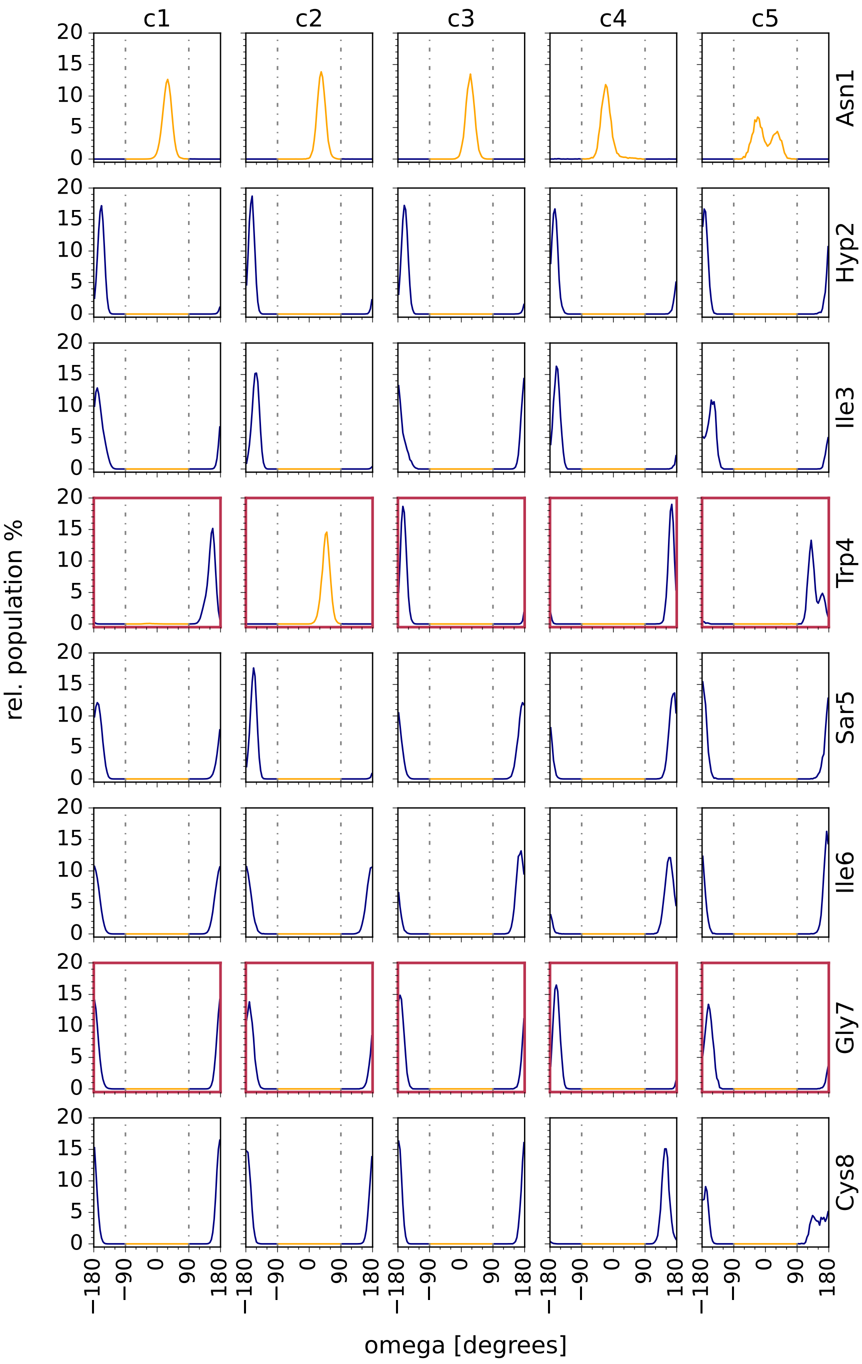

**Supplementary Figure 16.** Distributions of the $\omega$-torsion angles for the different residues in $M_{\text{ansa}}$-Gly5Sar-amanullin. Shown are the distributions over the clusters c1 - c5 that were identified for the sub data set 'tt' of the MD ensemble at 300 K. For the assignment of the trajectories to 'tt', please refer to Supplementary Table 5. All distributions were normalised to the size of the respective cluster. The $\omega$-torsion angles were defined according to IUPAC nomenclature (see 3.2.2). To discriminate the 'cis' configuration (orange) from the 'trans' configuration (blue), the threshold $\pm 90°$ was used (grey line). For $\omega$(Trp4) and $\omega$(Gly7), based on which the 'tt' sub set was defined, the plots are highlighted in dark red.

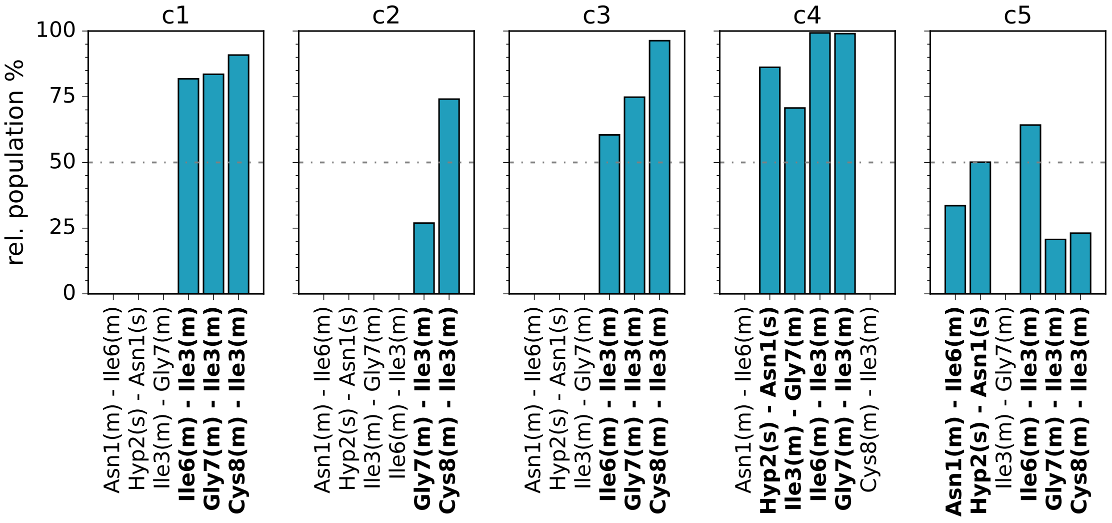

**Supplementary Figure 17.** Hydrogen bond populations ('bars') for the clusters c1 - c5 of the sub data set 'tt' of the $M_{\text{ansa}}$-Gly5Sar-amanullin MD ensemble at 300 K. For the assignment of the trajectories to 'tt', please refer to Supplementary Table 5. All hydrogen bond populations are normalised to the size of the respective cluster. Only hydrogen bonds are shown that occurred in at least 20% of the structures in at least one cluster (c1-c5). For each cluster, the hydrogen bonds with a relative occurrence ≥20% are additionally highlighted with bold font. All hydrogen bonds are labelled as 'donor - acceptor', and differentiated in main chain ('m', atoms: N, $H_N$, $C_\alpha$, $C_{CO}$ and $O_{CO}$) and side chain ('s', atoms: else) interactions. Please refer to 3.2.1 for further details on the computation of the hydrogen bonds.

##### 2.3.3 $P_{\text{ansa}}$-Gly5Sar-amanullin

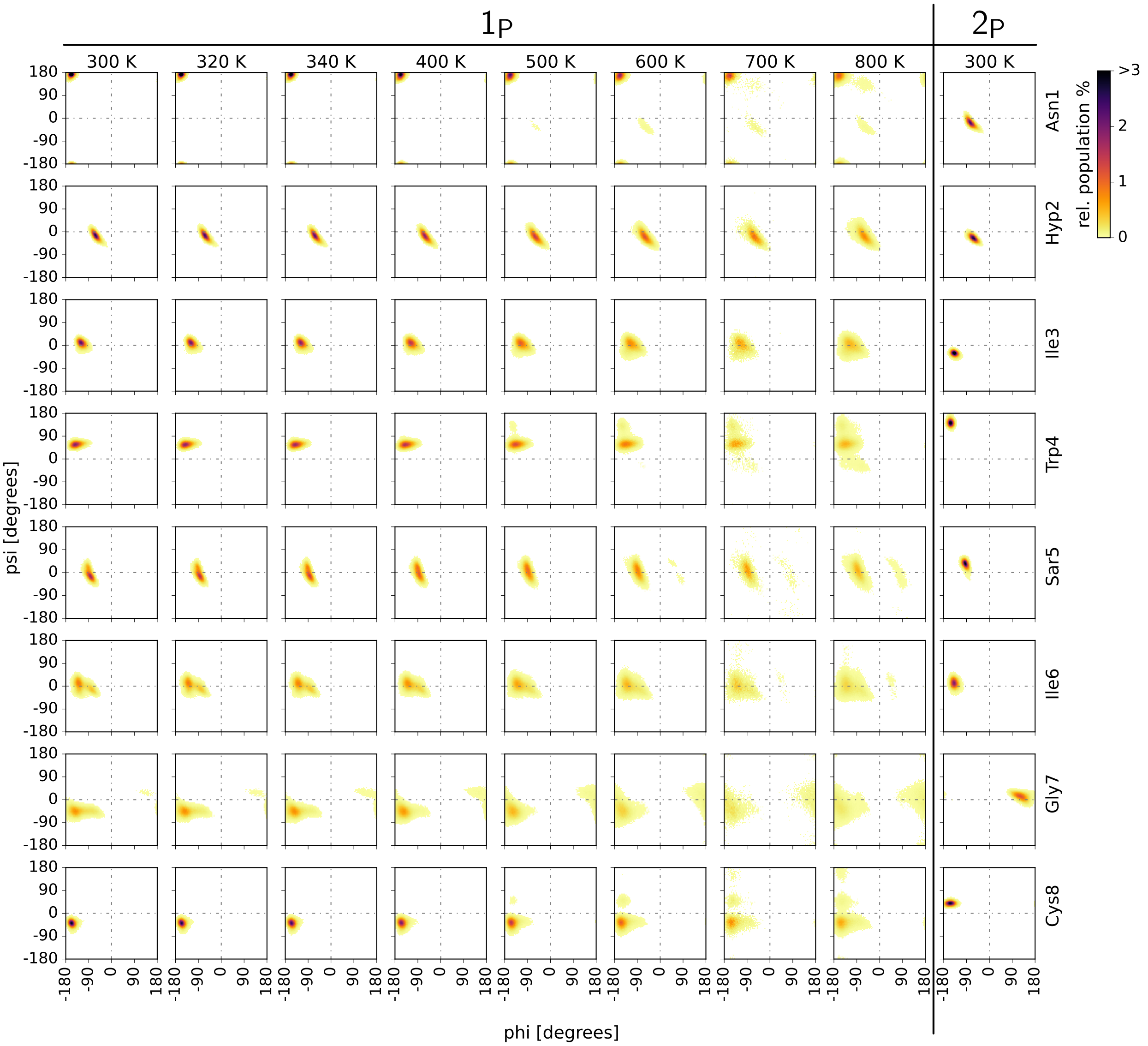

**Supplementary Figure 18.** Two-dimensional probability distributions of the $\phi$ and $\psi$ backbone torsion angles (Ramachandran plots) for the MD ensembles of $P_{\text{ansa}}$-Gly5Sar-amanullin at different temperatures. For an overview of the simulated data, please refer to Supplementary Table 10. The distributions shown represent the average distributions over all trajectories of the respective MD ensemble. For an overview of the amount of trajectories per ensemble, please refer to Supplementary Table 10. For the $P_{\text{ansa}}$-isomer, the simulations for $2_P$ were started from the 800 K MD ensemble of $1_P$. Therefore, $2_P$ is shown separately. Please refer to 3.1.4 and Supplementary Table 9.

**Supplementary Table 9.** Overview of the amount of trajectories that show an average, all-atom RMSD $\leq 0.1\,\mathrm{nm}$ to either their respective starting structure (‘A’) or to the highest-probability structure $1_{\mathrm{P}}$ of the $\mathrm{P_{ansa}}$-Gly5Sar-amanullin (‘B’). The average for the RSMDs was calculated referring to the simulation time $t$ in each step (see 3.2.9). The total amount of structures considered in each step is shown, too ($N_{\mathrm{s}}$).

| Step | $t$ | A | B | $N_{\mathrm{s}}$ |
|---|---|---|---|---|
| EM | – | 120 | 0 | 120 |
| NVT | 100 ps | 120 | 0 | 120 |
| NPT | 100 ps | 120 | 0 | 120 |
| RUN1 | 100 ps | 71 | 3 | 120 |
| RUN2 | 400 ps | 36 | 1 | 71 |
| RUN3 | 500 ps | 21 | 0 | 36 |
| RUN4 | 1 ns | 16 | 0 | 21 |
| RUN5 | 9 ns | 6 | 0 | 16 |
| RUN6 | 190 ns | 2 | 0 | 6 |

### 3 Computational methods

#### 3.1 MD simulations

##### 3.1.1 Overview of the simulation data

All MD simulations were performed with GROMACS[9–13] using the release versions 2019.4[14] and 2020.6.[15] In the following, the simulation setups for the MD simulations of the $P_{\mathrm{ansa}}$- and the $M_{\mathrm{ansa}}$-Gly5Sar-amnullin are described (see 3.1.3, 3.1.4, 3.1.5). For an overview of the final MD ensembles, please refer to Supplementary Table 10.

**Supplementary Table 10.** Overview of the production MD simulations performed to study the $P_{\mathrm{ansa}}$- and $M_{\mathrm{ansa}}$-isomers of Gly5Sar-amanullin. The considered ensembles, the simulated variants, the solvents, the simulation temperature $T$ and the simulation time $t_{\mathrm{sim}}$ are indicated.

| Ensemble | $T$ [K] | Solvent | $t_{\mathrm{sim}}$ [$\mu$s] | Sarcosine-variant |
|---|---|---|---|---|
| $NpT$ | 300 | DMSO | 24 | $M_{\mathrm{ansa}}$ |
| $NpT$ | 400 | DMSO | 24 | $M_{\mathrm{ansa}}$ |
| $NpT$ | 500 | DMSO | 24 | $M_{\mathrm{ansa}}$ |
| $NVT$ | 700 | DMSO | 24 | $M_{\mathrm{ansa}}$ |
| $NpT$ | 300 | DMSO | 12 | $P_{\mathrm{ansa}}$ ($1_P$) |
| $NpT$ | 320 | DMSO | 12 | $P_{\mathrm{ansa}}$ ($1_P$) |
| $NpT$ | 340 | DMSO | 12 | $P_{\mathrm{ansa}}$ ($1_P$) |
| $NpT$ | 400 | DMSO | 12 | $P_{\mathrm{ansa}}$ ($1_P$) |
| $NpT$ | 500 | DMSO | 12 | $P_{\mathrm{ansa}}$ ($1_P$) |
| $NVT$ | 600 | DMSO | 12 | $P_{\mathrm{ansa}}$ ($1_P$) |
| $NVT$ | 700 | DMSO | 12 | $P_{\mathrm{ansa}}$ ($1_P$) |
| $NVT$ | 800 | DMSO | 12 | $P_{\mathrm{ansa}}$ ($1_P$) |
| $NpT$ | 300 | DMSO | 50 | $P_{\mathrm{ansa}}$ ($2_P$) |

###### 3.1.2 Solvent boxes

Using the DMSO model parametrised by C. Caleman *et al.*,[16] three solvent boxes (A,B,C) were generated using 'gmx editconf' and 'gmx insert-molecules' of the GROMACS 2019.4 simulation software.[14] Solvent box A was equilibrated at 300 K, solvent box B was equilibrated at 350 K and solvent box C was equilibrated at 400 K, for 200 ps each.

###### 3.1.3 $P_{\text{ansa}}$-Gly5Sar-amanullin

###### 3.1.3.1 Structure construction

Following the structure of the *natural* amanullin,[2] a structure of the $P_{\text{ansa}}$-Gly5Sar-amanullin was built and energy minimised (UFF force field,[17] steepest-descent algorithm) with Avogadro 1.2.0.[18] Then, the structure was parametrised with ACPYPE[19] setting the molecule's charge $n = 0$ and referring to AMBER14SB force field[20] with $a =$'amber'.

###### 3.1.3.2 Generation of starting structures for MD simulations

The parametrised structure of $P_{\text{ansa}}$-Gly5Sar-amanullin was placed in each pre-equilibrated solvent box (A, B, C) using 'gmx solvate -cs *solvent box*' of the GROMACS 2019.4 simulation package.[14] All three boxes were energy minimised using the steepest-descent algorithm (emtol=1000 kJ/(mol·nm), nsteps=5000),[14] followed by $NVT$ and $NpT$ equilibrations at temperatures $T_A = 300$ K (box A), $T_B = 350$ K (box B) and $T_C = 400$ K (box C) for 400 ps ($NVT$) and 600 ps ($NpT$). In both equilibration steps, periodic-boundary conditions were applied in all directions and position restraints were used for the peptide structure ($f_x = f_y = f_z = 10^3$ kJ/(mol·nm$^2$)).

After equilibration, all three systems were simulated in the $NpT$-ensemble at temperatures $T_A = 300$ K (box A), $T_B = 350$ K (box B) and $T_C = 400$ K (box C) using leap-frog integration[21] with an integration time step d$t$ =2 fs. All covalent bonds were constrained using LINCS algorithm[22] ('all-bonds', iter=4, order=6). Periodic boundary conditions were applied on all three directions. The simulations were conducted using velocity-rescale thermostat[23] with coupling time $\tau_T$ =0.1 ps. The pressure was set to $p$ =1 bar using Parrinello-Rahman barostat[24] with coupling time $\tau_p$ =2 ps. For Van-der-Waals interactions, Verlet cut-off scheme[25] was applied with a cut-off radius of $r_{\mathrm{vdw}}$ =1 nm and an update of the neighborlist every nstlist=10 integration timesteps. For Coulomb interactions, Particle-Mesh-Ewald algorithm[26] was used (pme-order=6, Fourier grid spacing=0.12 nm, cut-off for short-range electrostatic interaction $r_{\mathrm{Coulomb}}$ =1 nm). The solute's coordinates were written to file every 1 ps over a simulation length of 0.1 $\mu$s.

Using 'gmx rms' of the GROMACS 2019.4 simulation package,[14] the all-atom RMSD of the peptide was calculated for each trajectory (A, B, C) after a least-squares fit on all $C_\alpha$ atoms. For each trajectory, the RMSD distribution was calculated using 'np.histogram()' of the numpy library[27] (range = (0,0.3) nm, bins = 100). Out of the four highest-populated bins of the RMSD distribution of each trajectory, one structure was randomly extracted yielding 4 structures per trajectory (in total 12). These 12 peptide structures were considered the starting structures for the MD simulations of the $P_{\mathrm{ansa}}$-Gly5Sar-amanullin. For the convenience of the reader, the origin of the starting structure of each trajectory is listed below: replicas 1-4 : trajectory A, replicas 5-8 : trajectory B, replicas 9-12 : trajectory C.

Using GROMACS 2019.4 release version,[14] each of the 12 peptide structures was placed in a box of identical size (cubic, $V \approx 51$ nm$^3$, 'gmx editconf -c'), solvated with the pre-equilibrated solvent box A (see 3.1.2, 'gmx solvate -cs') and energy minimised with steepest-descent algorithm (emtol = 1000 kJ/(mol·nm), nsteps=5000).

###### 3.1.3.3 Equilibrations

**[$T$ ≤500 K]** For the 12 solvated and energy-minimised starting structures, $NVT$ and $NpT$ equilibrations were performed at temperatures $T$ = 300 K, $T$ = 320 K, $T$ = 340 K, $T$ = 400 K and $T$ = 500 K for 400 ps ($NVT$) and 600 ps ($NpT$), respectively. Periodic boundary conditions were used in all directions and position restraints were applied on the peptides ($f_x = f_y = f_z = 10^3$ kJ/(mol·nm)).

**[$T$ >500 K]** For the temperatures higher than 500 K, $NVT$-equilibrations were performed in an iterative approach. The last frames of the $NVT$-equilibratio at temperature $T_1$ were used as starting structures for the $NVT$-equilibration at temperature $T_2$. The following pairs were considered ($T_1, T_2$): (500 K, 550 K), (550 K, 600 K), (600 K, 650 K), (650 K, 700 K), (700 K, 725 K), (725 K, 750 K), (750 K, 775 K) and (775 K, 800 K). All $NVT$-equilibrations were performed for 400 ps respectively.

###### 3.1.3.4 Production MD simulations

**[for all temperatures ≤500 K]** After equilibration, the 12 systems were propagated in the $NpT$-ensemble at the respective temperature using leap-frog integration[21] with an integration time step d$t$ =2 fs. LINCS algorithm[22] was used to constrain all bonds ('all-bonds', iter=4, order=6), and periodic boundary conditions were applied in all three directions. All simulations were performed with velocity-rescale thermostat[23] (coupling time $\tau_T$ =0.1 ps) and Parrinello-Rahman barostat[24] (coupling time $\tau_p$=2 ps, reference pressure $p$=1 bar). Van-der-Waals interactions were treated with Verlet cut-off scheme[25] (cut-off radius $r_{\mathrm{vdw}}$ =1 nm, update of the neighbourlist every nstlist=10 integration time steps). Coulomb interactions were treated with Particle-Mesh-Ewald algorithm (pme-order=6, Fourier grid spacing=0.12, cut-off for short-range electrostatic interactions $r_{\mathrm{Coulomb}}$ =1 nm). The solute's coordinates were written to file every 1 ps. For each trajectory, a simulation length of 1 $\mu$s was generated.

**[for temperatures $T$ with 500 K< $T$ ≤700 K]** After equilibration, the 12 systems were propagated in the $NVT$-ensemble at the respective temperature using the settings described for the MD simulations with $T$ ≤500 K except for the barostat, which was not used.

**[for $T$ = 800 K]** After equilibration, the 12 systems were propagated in the $NVT$-ensemble at $T = 800$ K using the settings described for the MD simulations with $T \leq 500$ K except for: LINCS algorithm[22] was only applied on bonds involving hydrogen atoms ('h-bonds', iter=4, order=6), and the barostat was not used.

###### 3.1.4 $P_{\text{ansa}}$-Gly5Sar-amanullin - further simulations

###### 3.1.4.1 Structures from the 800 K MD ensemble

Out of each 800 K trajectory of the MD simulations described in 3.1.3.4, ten structures were extracted (in total: 120) meeting the following criteria: The carbonyl-CO of Trp4 ('4m') is not involved in any hydrogen bond, and at least two structures per trajectory possess the hydrogen bond between the amide-NH of Trp4 and the carbonyl-CO of Cys8 ('4m-8m'). The so-extraced structures were placed and centred in a cubic simulation box ($V \approx 50$ nm$^3$) with GROMACS 2020.6 simulation package[15] using 'gmx editconf -c', and solvated with the pre-equilibrated solvent box A (see 3.1.2) using 'gmx solvate -cs *solvent box*'. The solvated systems were energy minimised using steepest-descent algorithm (emtol = 1000 kJ/(mol·nm), nsteps = 5000) with strengthened position restrains ($f_x = f_y = f_z = 10^5$ kJ/(mol·nm$^2$)).

###### 3.1.4.2 Equilibrations and pre-production RUNs

After energy minimisation, $NVT$ and $NpT$ equilibrations were performed at temperature $T$ = 300 K for 100 ps each. Periodic boundary conditions were used in all directions and default position restraints were applied on the peptides ($f_x = f_y = f_z = 10^3$ kJ/(mol·nm$^2$)). Then, the systems were simulated in the $NpT$-ensemble using the same parameters as described in 3.1.3.4 ($T \leq 500$ K), except for the simulation time. In total, six successive preruns were performed with simulation times $t_1$ =0.1 ns, $t_2$ =0.4 ns, $t_3$ =0.5 ns, $t_4$ =1 ns, $t_5$ =9 ns and $t_6$ =190 ns.

Thereby, the following approach was chosen: After each prerun, the all-atom RMSD was calculated with GROMACS 2020.6[15] using the the respective starting structure and the reference structure of $1_P$ (see 3.2.10). The RMSD calculations included a least-squares fit on all backbone atoms and all carbon atoms within one covalent bond of the backbone. For each system, the RMSDs were averaged over the respective simulation length. If the RMSD towards the starting structure was $\leq 0.1$ nm, the associated system was considered for the next prerun. Otherwise, it was discarded. After six preruns, only one structure met $\langle \text{RMSD} \rangle \leq 0.1$ nm towards its starting structure. For an overview of the assignment of the trajectories throughout the preruns, please refer to Supplementary Table 9.

###### 3.1.4.3 Production MD simulations

The structure that met the RMSD criterion described in 3.1.4.2 was simulated in the $NpT$-ensemble using the parameters and settings described in 3.1.3.4. In total, 50 trajectories with a simulation length of 1 $\mu$s were generated yielding a total simulation time of 50 $\mu$s (see Supplementary Table 10).

##### 3.1.5 $M_{\text{ansa}}$-Gly5Sar-amanullin

###### 3.1.5.1 Structure construction

The reference structures of the $P_{\text{ansa}}$-Gly5Sar-amanullin ($1_P$, $2_P$, see 3.2.10) were transformed to their $M$-ansameric form with Avogadro 1.2.0.[18] Thereby, the tryptathionine bridge was translocated to the opposite side of the macrolactam keeping the stereochemistry of the single amino acids intact. After construction, the structures were energy-minimised with Avogadro (UFF force field,[17] steepest-descent algorithm). If not stated otherwise, the parametrisation of the $P_{ansa}$-Gly5Sar-amanullin was used (see 3.1.3.1)

###### 3.1.5.2 Generation of starting structures for MD simulations

Using GROMACS 2020.6 simulation package, the so-prepared structures were placed in a cubic box (V≈ 50 $nm^3$, 'gmx editconf -c') and solvated with the pre-equilibrated solvent box A (see 3.1.2, 'gmx solvate -cs'). The solvated systems were energy minimised using steepest-descent algorithm (emtol = 1000 kJ/(mol·nm), nsteps=5000).

###### 3.1.5.3 MD simulations at $T$=500 K

For the solvated and energy-minimised structures (see 3.1.5.2), $NVT$- and $NpT$-equilibrations were performed at temperature $T$ = 500 K for 400 ps ($NVT$) and 600 ps ($NpT$), respectively. Periodic boundary conditions were used in all directions and position restraints were applied on the peptide ($f_x = f_y = f_z = 10^3$ kJ/(mol·nm)).

After equilibration, the systems were propagated using leap-frog integration[21] with an integration time step d$t$ = 2 fs. All covalent bonds were constrained using LINCS algorithm[22] ('all-bonds', iter=4, order=6). Periodic boundary conditions were applied in all three directions. The simulations were conducted in the $NpT$ ensemble at temperature $T$ = 500 K using velocity-rescale thermostat[23] with coupling time $\tau_T$ = 0.1 ps. The pressure was set to $p$ =1 bar using Parrinello-Rahman barostat[24] with coupling time $\tau_p$ =2 ps. For Van-der-Waals interactions, Verlet cut-off scheme[25] was applied with a cut-off radius of $r_{\mathrm{vdw}}$ =1 nm and a neighborlist-update every nstlist=10 integration timesteps. For Coulomb interactions, Particle-Mesh-Ewald algorithm[26] was used (pme-order=6, Fourier grid spacing=0.12 nm). The cut-off for short-range electrostatic interaction was set to $r_{\mathrm{Coulomb}}$ =1 nm. The solute's coordinates were saved every 1 ps. For each trajectory, a simulation length of 1 $\mu$s was generated. In total, 24 trajectories were generated yielding a total simulation time of 24 $\mu$s.

###### 3.1.5.4 MD simulations at $T$=400 K

The last frames of the 1 $\mu$s MD trajectories at 500 K (see 3.1.5.3) were equilibrated at temperature $T$ = 400 K for 400 ps ($NVT$) and 600 ps ($NpT$), respectively. Periodic boundary conditions were used in all directions and position restraints were applied on the peptide ($f_x = f_y = f_z = 10^3$ kJ/(mol·nm)). After equilibration, the systems were simulated in the $NpT$-ensemble at temperature $T$ = 400 K using the settings described for the production MD simulations at $T$ = 500 K. In total, 24 trajectories with a simulation length of 1 $\mu$s were generated yielding a total simulation time of 24 $\mu$s.

###### 3.1.5.5 MD simulations at $T$=300 K

The last frames of the 1 $\mu$s MD trajectories at 400 K (see 3.1.5.4) were equilibrated at temperature $T = 300$ K for 400 ps ($NVT$) and 600 ps ($NpT$), respectively. Periodic boundary conditions were used in all directions and position restraints were applied on the peptide ($f_x = f_y = f_z = 10^3$ kJ/(mol·nm)). After equilibration, the systems were simulated in the $NpT$-ensemble at temperature $T = 300$ K using the settings described for the production MD simulations at $T = 500$ K. In total, 24 trajectories with a simulation length of 1 $\mu$s were generated yielding a total simulation time of 24 $\mu$s.

###### 3.1.5.6 MD simulations at $T$=700 K

Starting from the last frames of the $NVT$-equilibration at $T = 500$ K, the ensuing equilibrations were performed in an iterative approach as described in 3.1.3.3. The last frames of the $NVT$ equilibrations at temperature $T_1$ were used as starting structures for $NVT$ equilibrations at temperature $T_2$. The following pairs were considered $(T_1, T_2)$: (500 K, 550 K), (550 K, 600 K), (600 K, 650 K), (650 K, 700 K). After equilibration, the systems were simulated in the $NVT$-ensemble at temperature $T = 700$ K using the settings described for the production MD simulations at $T = 500$ K except for: no barostat was used. In total, 24 trajectories with a simulation length of 1 $\mu$s were generated yielding a total simulation time of 24 $\mu$s.

#### 3.2 Analysis of MD simulations

Prior to the analyses described in the following, all MD trajectories were centred on the Gly5Sar-amanullin molecule using GROMACS 2020.6[15] ('gmx trjconv -pbc mol -center'). In addition, we applied a translational and rotational fit including a least-square fit on the $C_\alpha$-atoms ('-fit rot+trans'). If not stated otherwise, all operations indicated with 'gmx' were performed with GROMACS 2020.6 release version.[15]

##### 3.2.1 Hydrogen bonds

###### 3.2.1.1 Definitions

All atoms of the peptide were assigned to either the side chain ('s') or the main chain ('m') of the included residues using 'gmx make_ndx'. For the assignment, we referred to the definitions in GROMACS:[15] N, $H_N$, $C_\alpha$, $C_{CO}$ and $O_{CO}$ were considered 'main chain' atoms. All other atoms were assigned to the 'side chain' of the respective residue. For Sar5, we introduced a third group called 'main chain II', to which we assigned all atoms of the methyl group that replaces the amide proton ($H_N$). The hydrogen bonds were calculated on atom level with 'gmx hbond'. Based on the atom assignment, the hydrogen bonds were then classified as side or main chain interactions of the involved residues.

###### 3.2.1.2 Population analyses for single hydrogen bonds

All hydrogen bonds were evaluated based on their relative occurrence $P_h$ over the total simulation length:

$$P_h = \frac{1}{N_{\text{rep}}} \sum_{i=1}^{N_{\text{rep}}} \left( \frac{1}{N_{f,i}} \sum_{j=1}^{N_{f,i}} p_{h,i,j} \right) \tag{1}$$

with $p_{h,i,j} = 1$, if the hydrogen bond $h$ is present in trajectory $i$ in frame $j$ and $p_{h,i,j} = 0$, otherwise. $N_{\text{rep}}$ denotes the number of considered trajectories and $N_{f,i}$ denotes the number of frames in trajectory $i$ (per trajectory: $10^6$). All hydrogen bonds with a relative occurrence greater or equal to 20% ($P_h \geq 20\%$) were considered significant. Please note, if the hydrogen bond populations were analysed for sub-data sets such as clusters or assignments to ($\omega$(Trp4),$\omega$(Gly7))-combinations (Supplementary Table 5), eq. 1 simplifies to:

$$P_{h,c} = \frac{1}{N_{f,c}} \sum_{j=1}^{N_{f,c}} p_{h,c,j}, \tag{2}$$

with $N_{f,c}$ as the number of frames in the respective sub-data set $c$.

###### 3.2.2 Backbone angles $(\phi, \psi, \omega)$

Time series of the backbone angles $(\phi, \psi, \omega)$ were extracted from the MD trajectories with 'gmx gangle -oall -g1 dihedral'. We referred to the IUPAC angle definitions listed below:

$\phi_i$: $\angle\,(\mathrm{C_{i-1}}, N_i, C_{\alpha,i}, C_i)$

$\psi_i$: $\angle\,(\mathrm{N_i}, C_{\alpha,i}, C_i, N_{i+1})$

$\omega_i$: $\angle\,(\mathrm{C_{\alpha,i}}, C_i, N_{i+1}, C_{\alpha,i+1})$

for any residue $i$ of the peptide of interest. For the backbone angles ($\phi$,$\psi$), two-dimensional probability distributions were calculated with 'np.histogram2d()' of the python library 'numpy'.[27] Thereby, each angle was discretised into 90 equally-sized bins. The resulting 90 x 90 grids are presented as 2D images (Ramachandran plots).

###### 3.2.3 $^3J_{\mathrm{H^N-H^\alpha}}$-coupling constants

Based on the $\phi$-angles extracted from the MD trajectories, $^3J_{\mathrm{H^N-H^\alpha}}$-values were determined using Karplus' equation:[28]

$$^3J_{\mathrm{H^N-H^\alpha}} = A \cdot \cos^2(\phi + \theta) + B \cdot \cos(\phi + \theta) + C, \tag{3}$$

with $A = 6.64$, $B = -1.43$, $C = 1.86$ and $\theta = -60°$ from Ref. 3. For the comparison to the NMR ensemble, the average and standard deviation over all MD trajectories were calculated for each angle $\phi$.

##### 3.2.4 Comparison of MD trajectories and experimental NOE distance restraints

Interatomic distances between protons $a$ and $b$ were calculated with 'gmx distance -oall'. With $\mathbf{r}_i(a,b) = (r_0, ..., r_{N-1})^{\mathrm{T}}$ as the vector containing the distances $r_f(a,b)$ for all $N$ frames $f$ of trajectory $i$, the following ensemble averages were calculated:[29,30]

$$R_{a,b}^{\mathrm{MD}} = \frac{1}{N_{\mathrm{rep}}} \sum_{i=1}^{N_{\mathrm{rep}}} \left\langle \mathbf{r}_i(a,b)^{-6} \right\rangle^{-1/6}, \tag{4}$$

with $N_{\mathrm{rep}}$ as the number of considered MD trajectories. For sub-data sets such as clusters, $(\langle \mathbf{r}(a,b)^{-6} \rangle)^{-1/6}$ was calculated over all frames of the respective cluster ($N_{\mathrm{rep}} = 1$).

Using the NOE-derived distances $R_{a,b}^{\mathrm{NMR}}$ from NMR experiments, the MD ensemble was evaluated by:

$$\Delta(R_{a,b}) = R_{a,b}^{\mathrm{MD}} - R_{a,b}^{\mathrm{NMR}}. \tag{5}$$

Any distance with $\Delta(R_{a,b}) > 0.1\,\mathrm{nm}$ was considered 'violated', i.e. in this distance, the MD ensemble deviates from the NMR ensemble more than the experimental uncertainty. For the classification of the types of NOE distances, the all protons were assigned to the main chain ('m') or side chain ('s') of the respective residue (see 3.2.1). For Sar5, the group 'main chain II' ('x') was introduced, which contains all hydrogen atoms from the methyl group bound to the amide-N. The NOE distances from NMR are listed in Supplementary Table 4. For the ensemble averages $R_{a,b}^{\mathrm{MD}}$, please refer to Supplementary Table 6.

##### 3.2.5 Discretisation with TICA

The backbone angles $(\phi, \psi, \omega)$ of the residues in $M_{\mathrm{ansa}}$-Gly5Sar-amanullin were used as basis for the discretisation of the MD trajectories. Prior to discretisation, all trajectories of the 'tt' sub data set were reduced in their time resolution by a factor 100 using 'gmx trjconv -skip 100'. All angles were translated to '[radians]' and expressed in $\sin(x)$ and $\cos(x)$ to maintain their periodicity. Based on the $(\sin(x), \cos(x))$ time series with reduced time resolution, time-independent component analysis (TICA) was performed using 'pyemma.coordinates.tica()' of the PyEMMA Python package 2.5.7.[31] The lag time $\tau$ was set to 20. The number of output dimensions was regulated by setting 'variance cut-off' to 0.9. The projection on the first four time-independent components ('TICs') was written to file. For the assignment of the trajectories to sub data set 'tt', please refer to Supplementary Table 5.

##### 3.2.6 Density-based clustering

The discretised MD trajectories with reduced time resolution were clustered in the space of the time-independent components (see 3.2.5) using the common-nearest-neighbour clustering algorithm.[32–34] We used the implementation called 'CommonNN', which is included in the python software 'scikit.learn'[35] in the module 'scikit-learn-extra' and available from GitHub (source-link, documentation-link). For the clustering, two clustering objects were created: One called 'original' that comprised all trajectories with reduced time information ($\approx 10^5$ data points), and one called 'train' that included every 10th data point of 'original'. The clustering was performed for 'train' and then mapped on 'original' using the 'predict' function. The final assignment of the individual frames to either specific clusters or noise was saved as 'label trajectories'.

The CommonNN algorithm identifies clusters based on the data point density of the data set. The clusters are determined by three parameters $R$, $N$ and $M$, which can be defined by the user. $R$ denotes the radius of a sphere around any given point in the data set. $N$ denotes the number of neighbours two data points $a$ and $b$ have to share at least to belong to the same cluster. In that regard, 'sharing' means that these neighbours are located in the intersection of the spheres with radius $R$ around the two points $a$ and $b$. $M$ denotes the minimal number of data points below which any given cluster is classified as noise. Data points not assigned to any cluster are classified as noise, too.

Please refer to the Supplementary Table 8 for an overview of the parameter combination

$(R, N, M)$. The ratio of noise points is reported alongside the parameters.

##### 3.2.7 core-set Markov state models

Based on the clustered MD trajectories with reduced time resolution, core-set Markov state models were constructed using the 'msm' package of the PyEMMA Python package 2.5.7.[31] The clusters identified by applying 'CommonNN' on the discretised sub data set 'tt' of the $M_{\text{ansa}}$-Gly5Sar-amanullin MD ensemble (see 3.2.6) were defined as 'cores'. The label trajectories, which contained the assignment of every single frame to either a specific cluster or noise were used as input for the MSM. Please note, for the usage of PyEMMA both the cores labels and the label trajectories had to be shifted by '-1', so that noise is labelled '-1' and the cluster numbering starts with '0'. Due to the reduced time information, a time step in the trajectories corresponds to 0.1 ns. To compute implied timescales, lag times from 2 to 400 steps (0.2 to 400 ns) were used. The final cs-MSM was constructed and analysed for a lag time $\tau$=250 steps, which corresponds to 25 ns.

##### 3.2.8 Solvent-accessible surface area

Using GROMACS ('gmx sasa'), the solvent-accessible surface area (SASA) was calculated for the entire molecule (total), the hydrophobic atoms and the hydrophilic atoms. To achieve this, all atoms were classified as 'hydrophobic' if their charge (elementary charge [e]) is between 0.2 e and -0.2 e, and 'hydrophilic' otherwise. The SASA for the hydrophobic and hydrophilic atoms can then be computed by using the 'output'-option in 'gmx sasa':

```
gmx sasa -output '"Hydrophobic" group "AMA" and charge {-0.2 to 0.2};
         "Hydrophilic" group "AMA" and not charge {-0.2 to 0.2};'
```

with 'AMA' as the name for the simulated peptide. To trace back the assignment of the atoms by GROMACS, 'gmx select' is strongly advised:

```
gmx select -select '"Hydrophobic" ... '
```

##### 3.2.9 RMSD calculations

RMSD calculations were performed with 'gmx rms' including a least-squares fit on (a) all atoms or (b) all backbone atoms and carbon atoms within 1 covalent bond from the backbone. The selected group is always indicated.

##### 3.2.10 Reference structure of $P_{\mathrm{ansa}}$-Gly5Sar-amanullin

Based on the hydrogen bond populations $P_h$ introduced in 3.2.1, the hydrogen bonds with a relative occurrence $\geq 20\%$ over all trajectories of the respective $P_{\mathrm{ansa}}$-Gly5Sar-amanullin 300 K-MD ensembles ($1_P$, $2_P$) were determined. The combinations of these hydrogen bonds were evaluated as follows: Let $\mathbf{p}_{h,i}$ be the vector representation of the time series of hydrogen bond $h$ in MD trajectory $i$. Considering all $n_h$ hydrogen bonds with a relative occurrence $\geq 20\%$, a vector of combination labels $\mathbf{X}_i$ was calculated for each trajectory $i$ with:

$$\mathbf{X}_i^{\mathrm{T}} = \sum_{h=0}^{n_h} 2^k \cdot \mathbf{p}_{h,i}^{\mathrm{T}}, \tag{6}$$

where $\mathbf{X}_i$ denotes the trajectory $i$ expressed in combination labels from 0 to $C = \sum_{h=0}^{n_h} 2^h$. The hydrogen bond combination with highest probability over all trajectories was considered most important. All associated frames were extracted from the respective $P_{\mathrm{ansa}}$-Gly5Sar-amanullin 300 K-MD ensembles ($1_P$, $2_P$) and saved to file as sub data sets. The first frame of these sub data set were considered as reference structures for the RMSD calculations (see 3.1.4) and structure generations of the $M$-ansamers (see 3.1.5.1).

### 4 Experimental methods

#### 4.1 HPLC-MS

Preparative HPLC was carried out on a 1260 Infinity (Agilent Technologies, Waldbronn, Germany) HPLC system with a polymeric reversed phase column (PLRP-S 100A) 300 x 50 mm, particle size 10 $\mu$m, Agilent Technologies, Waldbronn, Germany).

HPLC-MS: HPLC-HRMS spectra were recorded on a QTrap LTQ XL (Thermo Fisher Scientific, Waltham, Massachusetts, USA) hyphenated to an Agilent 1200 Series HPLC-System (Agilent Technologies, Waldbronn, Germany) equipped with a C18 column (50 x 2 mm, particle size 3 $\mu$m). HPLC-HRMS chromatograms were obtained with a solvent gradient of 0.1% formic acid in water (Solvent A) and 0.1% formic acid in acetonitrile (Solvent B).

The solvent gradients are shown below:
Gradient: 0-10 min 10%-50% B, 10-13 min 100% B, 13-16 min 20% B.

#### 4.2 CD spectroscopy

Circular dichroism spectra were measured on a J-815 CD spectrometer (Jasco, Groß-Umstadt, Germany). The lyophilized compound was dissolved in $H_2O$ to reach a concentration of 75µM. Far-UV spectra were acquired at 20 °C between 190-300 nm with a path length of 0.1 cm. The bandwidth was set to 1 nm at a continuous scanning speed of 50 nm/min in 5 accumulations. The data pitch was set to 0.1 nm. The spectra were processed with Spectra Manager (JASCO) and the mean residue ellipticity (MRE) was calculated as follows:

$$\mathrm{MRE} = \frac{\theta \cdot 0.1}{l \cdot c \cdot n}, \tag{7}$$

where $\theta$ is the ellipticity and $l$, $c$, $n$ denote path length, molar concentration and number of amino acids.

#### 4.3 NMR spectroscopy

##### 4.3.1 Setup

NMR spectra were recorded at 298 K using the following spectrometers: Bruker Avance-II 400 MHz, Bruker Avance-III 500 MHz or Bruker Avance III 700 MHz (Bruker, Karlsruhe, Germany). The chemical shifts are reported in ppm using the residual solvent peak as an internal reference (DMSO-$d_6$). Multiplicity (br. s = broad singlet, s = singlet, d = doublet, dd = doublet of doublet, t = triplet, q = quartet, m = multiplet) and coupling constants ($[J]$ = Hz) are quoted where possible.

##### 4.3.2 NMR assignment

To obtain resonance assignments $M_{\text{ansa}}$-Gly5Sar-amanullin was dissolved in deuterated DMSO-d7 (approx. 0.01M). TOCSY, COSY and NOESY spectra were recorded on a Bruker Avance III 700 MHz spectrometer with a TXI 5 mm probe. Standard Bruker pulse programs were used and all spectra were acquired at 298 K. Residual solvent methyl peaks (DMSO-d7 $\delta$ = 2.50 ppm for $^{1}$H and $\delta$ = 39.5 ppm for $^{13}$C) were used for chemical shift referencing. 2D homonuclear spectra were measured with acquisition times of 70 ms and 18 ms for the direct and indirect dimensions, respectively. TOCSY and NOESY spectra were accumulated with 32 scans and COSY spectra with 8 scans. The TOCSY and NOESY mixing times were set to 100 ms and 300 ms, respectively. Natural abundance $^{13}$C-HSQC spectra were measured with 140 scans and acquisition times of 14 ms and 120 ms for the direct and indirect dimensions. For calculation of the exchange coefficient, EXSY spectra were acquired at mixing times of 0 ms, 250 ms and 500 ms. The rate constants were calculated using EXSYCalc 1.0.

The spectra were processed and analysed using TopSpin 3.5 (Bruker) and CcpNmr 2.3.1 (The CCPN data model for NMR spectroscopy: development of a software pipeline).[36] The NOE correlations were manually assigned in CcpNmr 2.3.1.

##### 4.3.3 Variable temperature NMR (VT-NMR)

$^{1}$H-NMR spectra were acquired from 300 K to 350 K in 10 K increments in DMSO-d6. Temperature coefficients were obtained from the correlation between amide chemical shifts and temperature, and are categorized as the following: $\Delta\delta$HN/$\Delta$T with values less than -4.6 ppb/K indicate solvent-exposed NHs. Intermediate values from –4.6 to –3.0 ppb/K indicate intermediate shielding and potentially weak or strained hydrogen bonding. $\Delta\delta$HN/$\Delta$T values greater than –3.0 ppb/K indicate that the amides are highly shielded and potentially strongly hydrogen bound.

(6) Pardi, A.; Billeter, M.; Wüthrich, K. Calibration of the angular dependence of the amide

proton-C$\alpha$ proton coupling constants, 3JHN$\alpha$, in a globular protein: Use of 3JHN$\alpha$ for identification of helical secondary structure. *J. Mol. Biol.* **1984**, *180*, 741–751.

(12) Páll, S.; Abraham, M. J.; Kutzner, C.; Hess, B.; Lindahl, E. Tackling exascale software challenges in molecular dynamics simulations with GROMACS. International conference on exascale applications and software. pp 3–27.

(13) Abraham, M. J.; Murtola, T.; Schulz, R.; Páll, S.; Smith, J. C.; Hess, B.; Lindahl, E. GROMACS: High performance molecular simulations through multi-level parallelism from laptops to supercomputers. *SoftwareX* **2015**, *1*, 19–25.

(14) Abraham, M. J.; van der Spoel, D.; Lindahl, E.; Hess, B.; GROMACS development team, GROMACS 2019.4 Manual. 2019.

(15) Abraham, M. J.; van der Spoel, D.; Lindahl, E.; Hess, B.; GROMACS development team, GROMACS 2020.6 Manual. 2021.
